# Anti-diabetic drug binding site in K_ATP_ channels revealed by Cryo-EM

**DOI:** 10.1101/172908

**Authors:** Gregory M. Martin, Balamurugan Kandasamy, Frank DiMaio, Craig Yoshioka, Show-Ling Shyng

## Abstract

Sulfonylureas are anti-diabetic medications that act by inhibiting pancreatic K_ATP_ channels composed of SUR1 and Kir6.2. The mechanism by which these drugs interact with and inhibit the channel has been extensively investigated, yet it remains unclear where the drug binding pocket resides. Here, we present a cryo-EM structure of the channel bound to a high-affinity sulfonylurea drug glibenclamide and ATP at 3.8Å resolution, which reveals in unprecedented details of the ATP and glibenclamide binding sites. Importantly, the structure shows for the first time that glibenclamide is lodged in the transmembrane bundle of the SUR1-ABC core connected to the first nucleotide binding domain near the inner leaflet of the lipid bilayer. Mutation of residues predicted to interact with glibenclamide in our model led to reduced sensitivity to glibenclamide. Our structure provides novel mechanistic insights of how sulfonylureas and ATP interact with the K_ATP_ channel complex to inhibit channel activity.

## Introduction

ATP-sensitive potassium (K_ATP_) channels are unique hetero-octameric complexes each composed of four inwardly rectifying Kir6 channel subunits and four sulfonylurea receptor (SUR) subunits belonging to the ATP binding cassette (ABC) transporter protein family (Aguilar-Bryan and Bryan, 1999; Nichols, 2006). In pancreatic β-cells, K_ATP_ channels formed by Kir6.2 and SUR1 are gated by intracellular ATP and ADP, with ATP inhibiting channel activity while Mg^2+^-complexed ATP and ADP stimulating channel activity (Aguilar-Bryan and Bryan, 1999; Ashcroft, 2007). During glucose stimulation, the intracellular ATP to ADP ratio increases following glucose metabolism, which favors channel closure by ATP, resulting in membrane depolarization, Ca^2+^ influx, and exocytosis of insulin granules. In this way, K_ATP_ channels are able to control insulin secretion according to blood glucose levels. Mutations that disrupt channel function are known to cause a spectrum of insulin secretion disorders (Ashcroft, 2005; Koster et al., 2005). Specifically, loss-of-function mutations result in congenital hyperinsulinism, whereas gain-of-function mutations lead to transient or permanent neonatal diabetes (Ashcroft, 2005). The pivotal role of K_ATP_ channels in insulin secretion regulation makes them an important drug target.

Discovered in the 1940s, sulfonylureas have been a mainstay of type 2 diabetes therapy for more than half a century (Sola et al., 2015). The medical importance of this class of drugs has led to its evolution into several generations of agents, including first-generation sulfonylureas such as tolbutamide and second-generation agents such as the high-affinity sulfonylurea glibenclamide (GBC) (Gribble and Reimann, 2003; Sola et al., 2015). All sulfonylureas stimulate insulin secretion to reduce plasma glucose levels by blocking the activity of β-cell K_ATP_ channels (Gribble and Reimann, 2003). More recently, they have also become the primary pharmacotherapy for neonatal diabetes patients carrying gain-of-function K_ATP_ channel mutations (Aguilar-Bryan and Bryan, 2008; Ashcroft, 2007; Sagen et al., 2004). Despite their clinical importance and decades of research, how sulfonylureas interact with and block K_ATP_ channel activity remains poorly understood.

To begin to address the structural mechanisms by which ATP and sulfonylureas such as GBC inhibit K_ATP_ channels to stimulate insulin secretion, we carried out single particle cryo-EM and determined the structure of the β-cell K_ATP_ channel complex in the presence of ATP and GBC (Martin et al., 2017). While our initial structure at a resolution of 5.7Å revealed the overall architecture of the channel and location of the ATP molecule, it was unable to clearly define the GBC binding site and the atomic details associated with ATP binding. In this study, we improved the resolution of the K_ATP_ channel structure bound to GBC and ATP to 3.8Å. The higher resolution structure not only clearly defines the GBC and ATP binding pockets but also provides novel insights into the mechanisms of channel inhibition by ATP and GBC.

## Results

### Structure determination

To obtain a structure of K_ATP_ channels bound to GBC and ATP, channels comprising a rat Kir6.2 and FLAG-tagged hamster SUR1 (96 and 95% sequence identity to human sequences, respectively) were expressed in rat insulinoma INS-1 832/13 cells (Hohmeier et al., 2000), affinity purified, and imaged in the presence of 1mM ATP (no Mg^2+^) and 1µM GBC, as described previously (Martin et al., 2017). To improve resolution, we adjusted sample and grid preparation parameters (see Materials and Methods) to optimize ice thickness and particle orientation distributions, which both increased the overall quality and quantity of single particles.

3D classification in RELION yielded one four-fold symmetric class, which reached an overall resolution of 3.8Å after refinement (Fig. 1-figure supplements 1; Table 1). The local resolution, as estimated by Bsoft, varied from 3.2Å in the Kir6.2 transmembrane domain (TMD) to 5.2Å in the SUR1 nucleotide binding domains (NBDs) (Fig. 1-figure supplements 2). Overall, the map displays excellent connectivity to allow for model building (Fig.1). We have constructed a full atomic model for all of Kir6.2 minus disordered N- and C-termini, and for TMD0 of SUR1, as this part of the map was almost entirely at or below 3.7Å, with clear side-chain density for most residues. The ABC core of SUR1 displayed greater variability in resolution: the inner helices (relative to Kir6.2/TMD0) were also very well resolved (between 3.5 and 4Å resolution) to permit nearly complete atomic model building, while the most exposed helices (TMs 9, 10, 12, and 13) showed signs of flexibility, and were only built as polyalanine chains. This was also the case for the NBDs, for which we only refined our previously-deposited NBD models as rigid bodies (see Materials and Methods).

**Table 1.**
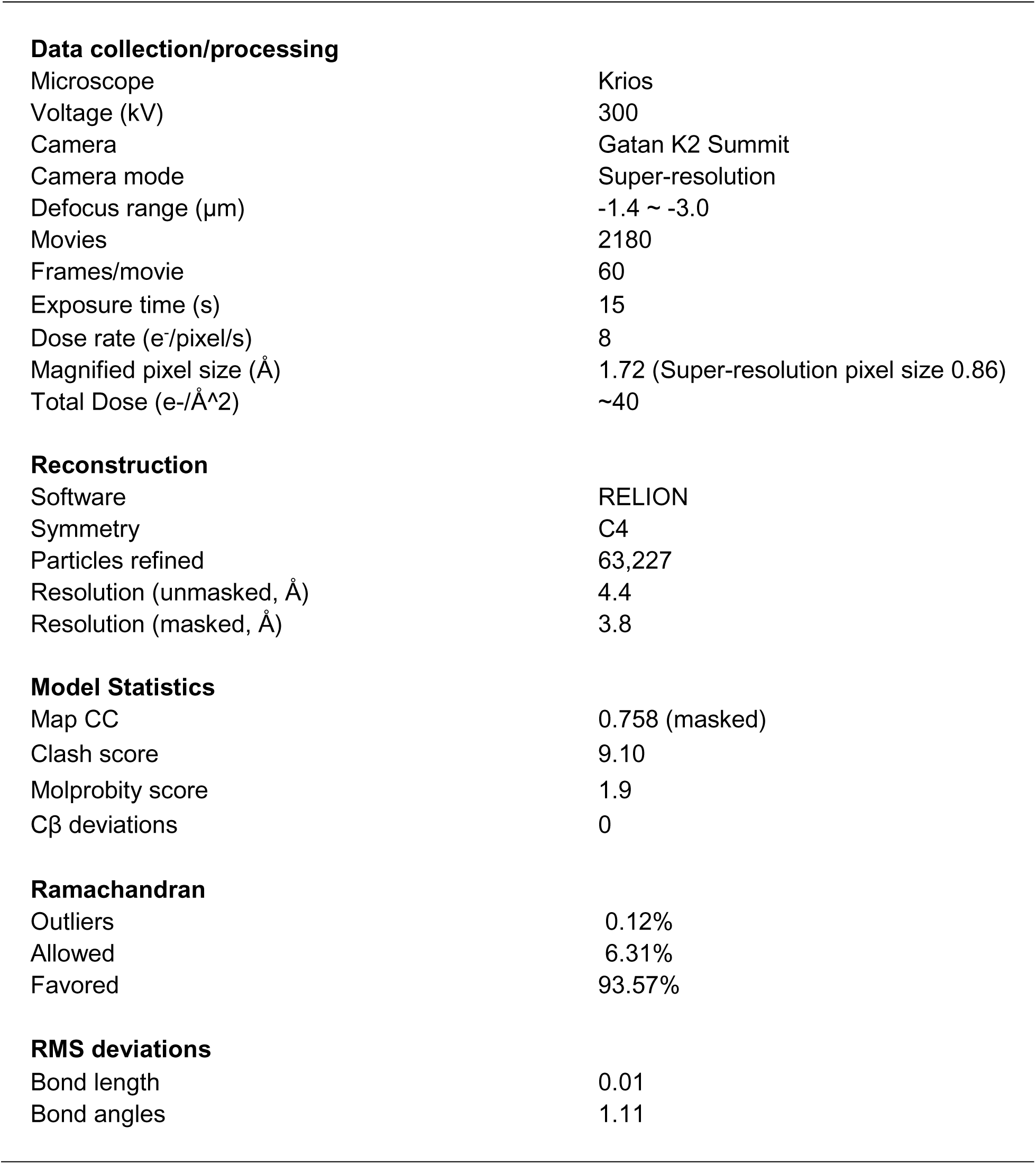
Statistics of cryo-EM data collection, 3D reconstruction and model building.

**Data collection/processing**
|  |  |
| --- | --- |
| Microscope | Krios |
| Voltage (kV) | 300 |
| Camera | Gatan K2 Summit |
| Camera mode | Super-resolution |
| Defocus range ($\mu\text{m}$) | -1.4 ~ -3.0 |
| Movies | 2180 |
| Frames/movie | 60 |
| Exposure time (s) | 15 |
| Dose rate ($\text{e}^-/\text{pixel}/\text{s}$) | 8 |
| Magnified pixel size ($\text{\AA}$) | 1.72 (Super-resolution pixel size 0.86) |
| Total Dose ($\text{e}^-/\text{\AA}^2$) | ~40 |
| **Reconstruction** |  |
| Software | RELION |
| Symmetry | C4 |
| Particles refined | 63,227 |
| Resolution (unmasked, $\text{\AA}$) | 4.4 |
| Resolution (masked, $\text{\AA}$) | 3.8 |
| **Model Statistics** |  |
| Map CC | 0.758 (masked) |
| Clash score | 9.10 |
| Molprobity score | 1.9 |
| Cβ deviations | 0 |
| **Ramachandran** |  |
| Outliers | 0.12% |
| Allowed | 6.31% |
| Favored | 93.57% |
| **RMS deviations** |  |
| Bond length | 0.01 |
| Bond angles | 1.11 |

**Fig. 1.**
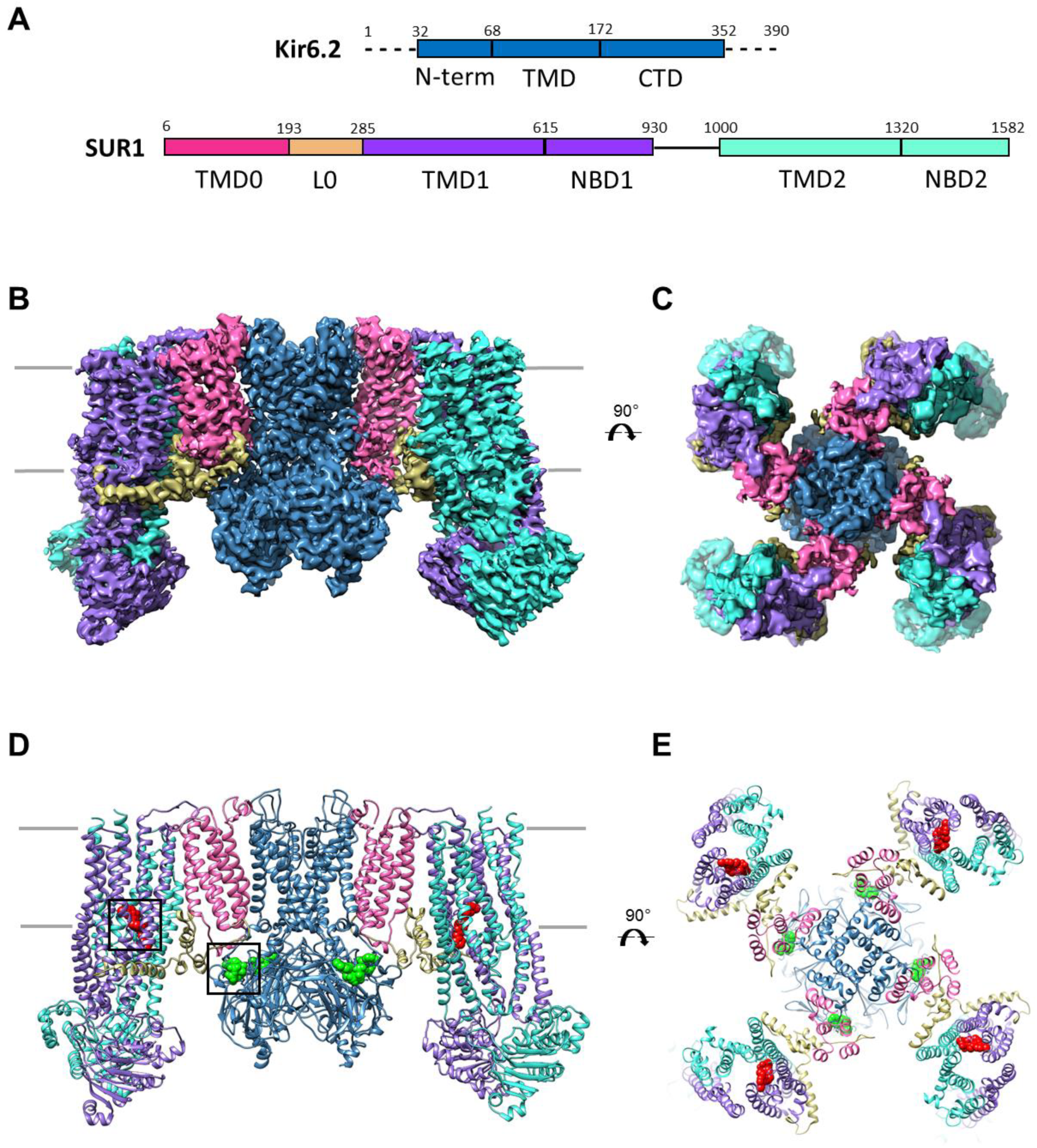
Overall structure of the K_ATP_ channel bound to ATP and GBC. (**A**) Linear sequence diagram for the Kir6.2 and SUR1 polypeptides, with primary domains colored to match the panels below. Numbers indicate residue number at the beginning and end of each domain. (**B**) Cryo-EM density map of the K_ATP_ channel complex at 3.8Å resolution, viewed from the side. Gray bars indicate approximate position of the bilayer. (**C)** View of map from extracellular side. (**D**). Structural model of the complex, with ligands ATP (green) and GBC (red) in boxes. (**E**) View of the model from the extracellular side.

### Structural overview

The K_ATP_ channel is built around a tetrameric Kir6.2 core with each subunit in complex with one SUR1 (Fig.1B-E), as observed previously (Martin et al., 2017). Each Kir6.2 has the typical Kir channel architecture of an N-terminal cytoplasmic domain, a TMD consisting of two TMs termed M1 and M2 interspersed by a pore loop and selectivity filter, and a “tether” helix that links the TMD to the larger C-terminal cytoplasmic domain (CTD) (Fig.2A, Fig.2-figure supplement 1). In our new structure, constrictions in the selectivity filter (T130), bundle crossing (F168), and the G-loop (G295, I296) are clearly seen (Fig.2B, C) to indicate a closed pore, as expected given the channel is bound to ATP.

**Fig. 2.**
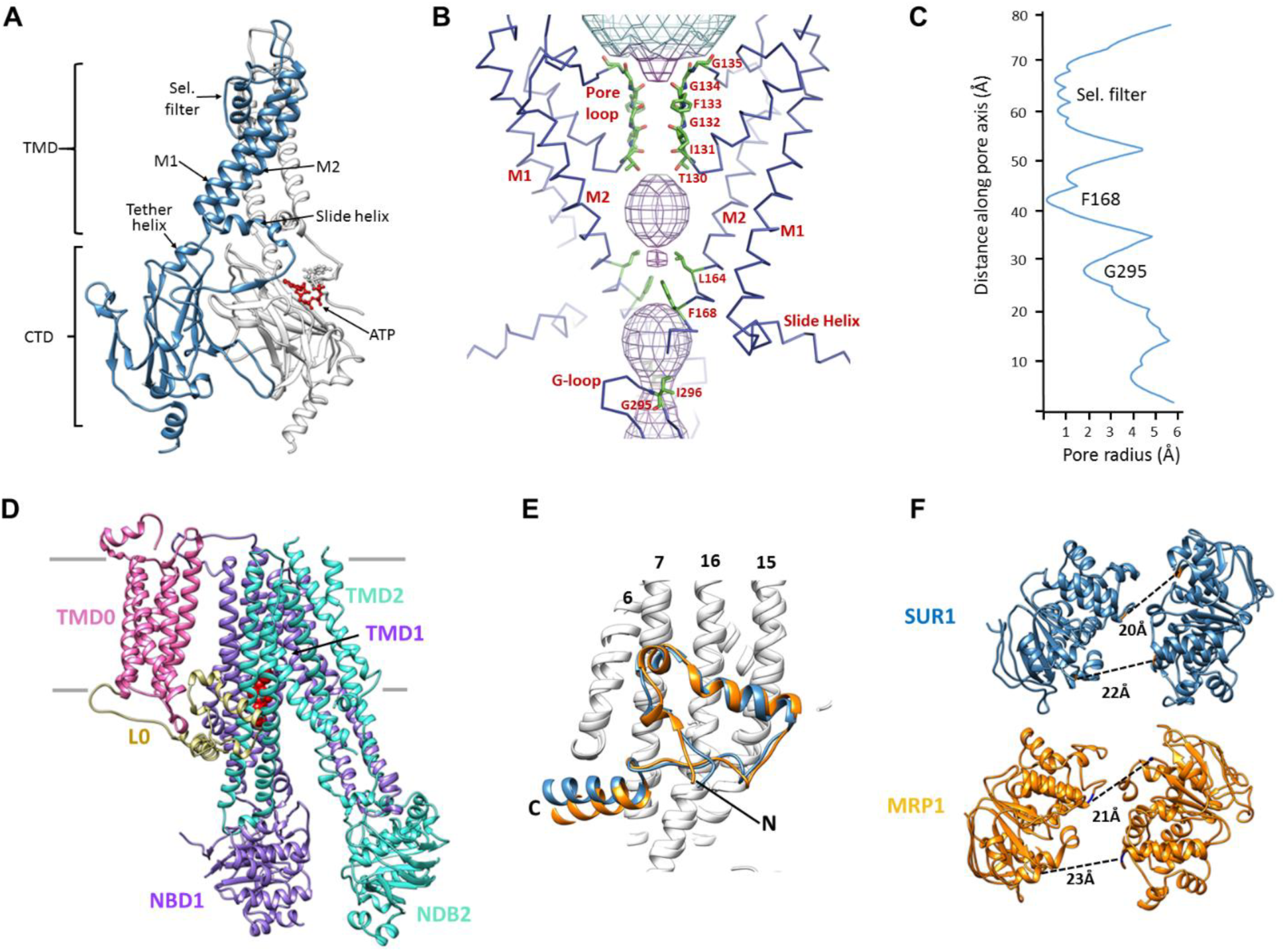
Structural highlights of Kir6.2 and SUR1. (**A**) Two subunits of the Kir6.2 tetramer, one colored in blue and one in white, highlighting the conserved Kir channel structural features. Note the ATP-binding site is at the interface of the cytoplasmic N- and C-terminal domains of adjacent subunits. (**B**) Close-up of the Kir6.2 pore, showing solvent-accessible volume as a mesh. The two primary gates are 1) the helix bundle-crossing (HBC), formed by the confluence of the M2 helices at F168; 2) the G-loop, formed at the apex of the CTD by G295 and I296. (**C**) Plot of pore radius as a function of length along pore axis. (**D**) Structure of SUR1 in inward-facing conformation, indicating overall domain organization. Note clear separation of NBDs. (**E**) Structural conservation of L0 with the lasso domain observed in MRP1. Full structures of SUR1 (blue) and leukotriene C4-bound MRP1 (orange) minus TMD0 were used for structural alignment. (**F**) Separation (Cα to Cα, indicated by the dashed line) between Walker A and signature motif in NBD1 (left) and NBD2 (right) (G716::S1483 and S831::G1382 in SUR1, G681::S1430 and S769::G1329 in MRP1).

SUR1 is one of only a handful of ABC transporters which possess an N-terminal transmembrane domain, TMD0, in addition to an ABC core structure comprising two TMDs of 6 helices each and two cytosolic NBDs (Tusnady et al., 2006; Wilkens, 2015) (Fig.2D). In the structure, TMD0 is a well-resolved 5-TM bundle (Fig.2D, Fig.2-figure supplement 2). A long intracellular loop L0 which tethers TMD0 to the ABC core is found to contain both cytosolic and amphipathic domains (Fig.2D, Fig.2-figure supplement 2). The C-terminal 2/3 of L0 is homologous to the “lasso motif” observed in CFTR and MRP1 (Johnson and Chen, 2017; Liu et al., 2017; Zhang and Chen, 2016), and indeed, the structures are very similar (Fig.2E). SUR1 is found in an “inward-facing” conformation, with NBDs clearly separated and the vestibule formed by TMD1/TMD2 open towards the cytoplasm. As we noted previously, the two TMD-NBDs show a ~15° rotation and ~10Å horizontal translation relative to each other (Fig.2F). This lack of symmetry is also seen in recently reported CFTR and MRP1 inward-facing structures. The separation between the two NBDs in our structure is similar to that seen in MRP1 bound to its substrate leukotriene C4 (see Fig.2F). Like SUR1 in which only NBD2 is capable of hydrolyzing ATP while NBD1 harbors a degenerate ATPase site, CFTR and MRP1 also have two asymmetric NBDs, suggesting the relative rotation and translation between the two TMD-NBD halves may be a common characteristic of ABC transporters with asymmetric NBDs.

### Unique molecular interactions between SUR1 and Kir6.2

Among all Kir channels, Kir6.1/Kir6.2 are the only members known to couple to an ABC transporter; and among all ABC transporters, SUR1/SUR2 are the only ones known to couple to an ion channel. These proteins are also unique in that they are co-dependent for both expression and function (Inagaki et al., 1995; Zerangue et al., 1999). How SUR1 and Kir6.2 achieve this unique regulation has been a long standing question in the field.

In the structure, we find a series of hydrophobic and polar interactions mediated exclusively by TMD0 and L0 with Kir6.2 (Fig. 3). The extracellular N-terminus of SUR1 closely contacts the turret and pore loop of Kir6.2 (Fig.3A-D), while TM1 of TMD0 and the M1 helix of Kir6.2 form a series of hydrophobic interactions running the length of the helices (Fig.3E). On the cytoplasmic side, the intracellular loops ICL1, ICL2, and the N-terminal portion of L0, prior to the “lasso motif,” cluster around the Kir6.2 N-terminus which harbors the slide helix and key ATP-binding residues and also forms part of an intersubunit β-sheet (Fig.2B).

**Fig. 3.**
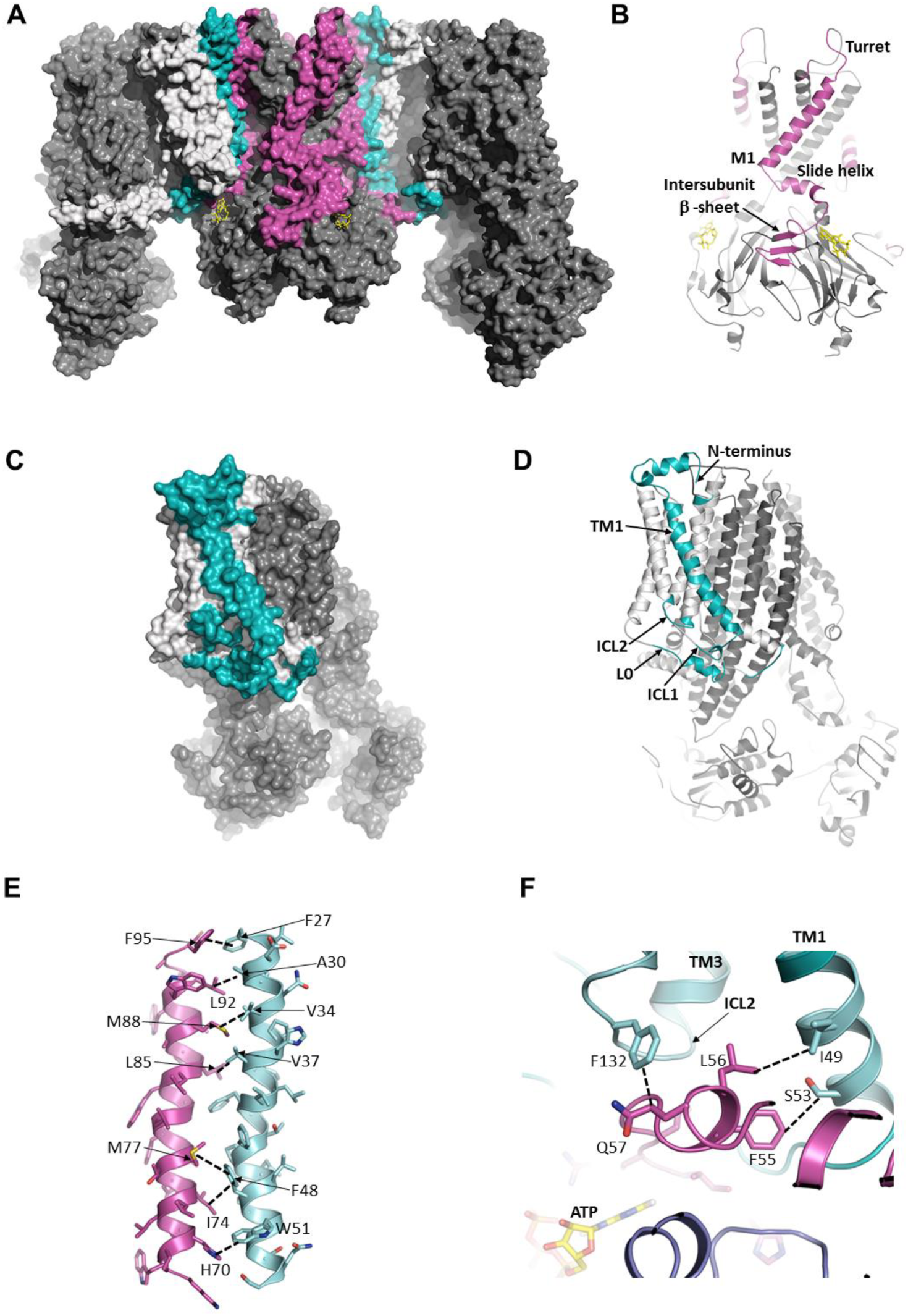
The interface between SUR1 and Kir6.2. (**A**) Surface representation of the complex. SUR1-binding surface on Kir6.2 colored in magenta, and Kir6.2-binding surface on SUR1 is in cyan. TMD0/L0 is colored in light gray, and Kir6.2 and the ABC core of SUR1 are in dark gray. (**B**) Cartoon model of Kir6.2, with interface residues colored in magenta. The intersubunit β-sheet formed by β strands ANO shown in Fig.2-figure supplement 2. (**C** and **D**) Surface and cartoon models of SUR1, with interface residues in cyan. (**E**) Interface between M1 (Kir6.2; magenta) and TM1 (SUR1; cyan), highlighting key interactions. (**F**) Intersection of ICL2 (cyan) and N-terminus/slide helix (magenta), showing interaction between Q57 (Kir6.2) and F132 (SUR1). The dashed lines in this and subsequent figures indicate select van der Waals or electrostatic (H-bonding or charge-charge) between two residues to aid visualization.

Of all the Kir channel family members which display a high degree of sequence conservation, only Kir6.1 or Kir6.2 co-assemble with SUR proteins, raising the interesting question of what molecular interactions confer this specificity. While many residue pairs in the interface are conserved in either the Kir or ABC transporter family, a couple residue pairs are unique to both Kir6.2 and SUR1. Among these, H70 within the M1 helix of Kir6.2 forms an edge-to-face π-stacking interaction with W51 of TM1 of TMD0 (Fig.3E), and Q57 of the Kir6.2 slide helix contacts F132 of ICL2 (Fig.3F). F132 is a well-studied PNDM mutation which causes very high *P_o_* but also reduces physical interaction between TMD0 and Kir6.2 (Proks et al., 2007), supporting its role as a critical part of the interface. To our knowledge, mutational studies of Kir6.2 Q57 and H70, and SUR1 W51 have not been reported; it would be interesting to test the role of these residues in channel assembly and function in the future.

### The ATP binding site

Non-hydrolytic binding of intracellular ATP to the cytoplasmic domains of Kir6.2 induces rapid and reversible closure of the pore (Nichols, 2006). We have previously reported the location of the ATP-binding site at the interface of the cytoplasmic N- and C-terminal domains from two adjacent subunits (Martin et al., 2017), giving 4 equivalent sites for the Kir6.2 tetramer. In the current map, there is strong electron density for the ATP as well as surrounding residues (Fig.4., Fig.4-figure supplement 1), allowing for detailed analysis of the mode of ATP binding as well as the mechanism of inhibition.

**Fig. 4.**
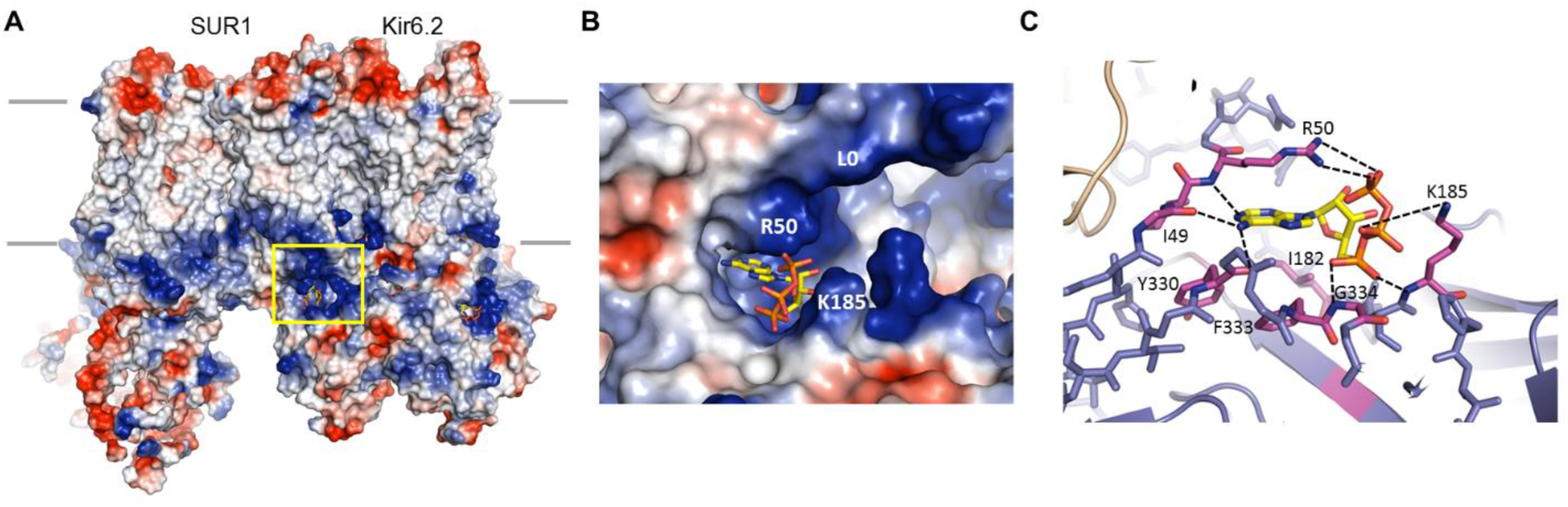
The ATP binding pocket. (**A**) Surface representation of a Kir6.2 tetramer in complex with one SUR1, colored by Coulombic surface potential. ATP pocket is boxed in yellow. (**B**) Close-up of ATP binding pocket boxed in (**A**). Note close proximity of L0 to the pocket on Kir6.2. (**C**) Atomic interactions within ATP-binding pocket, with residues directly interacting with ATP colored in magenta.

In the structure, the ATP is directly below the inner membrane leaflet and is partially exposed to solvent (Fig.4). The bound ATP appears to adopt a conformation similar to that found in other non-canonical, Mg^2+^-independent ATP-binding sites, such as the P2X receptor (Hattori and Gouaux, 2012), in which the phosphate groups are folded towards the adenine ring (Fig.4B, C). This places the β- and γ-phosphates to interact with basic residues contributed by the N- and C-termini. The pocket itself is formed by the overlap of three distinct cytoplasmic structures: an N-terminal peptide (binding residues I49 and R50; subunit A) immediately before the Kir channel “slide helix”; a C-terminal β-sheet (I182 and K185; subunit B) immediately following the TMD-CTD tether helix (see Fig.2A, Fig.2-figure supplement 1); and a short, solvent-exposed helical segment (Y330, F333, G334; subunit B).

The α- phosphate of ATP is coordinated by the main-chain nitrogen of G334 and K185, while the β- and γ-phosphates are coordinated by side-chain nitrogens of K185 and R50, respectively. The ribose group is in close contact with the I182 and F333 side chains; the adenine ring stacks against the aliphatic portion of the R50 side chain as well as Y330, and is H-bonded to the main chain nitrogen of R50, and main chain carbonyls of I49 and Y330. The aforementioned residues have all been shown previously to reduce ATP inhibition when mutated to other amino acids (Antcliff et al., 2005; Cukras et al., 2002; Drain et al., 1998; Li et al., 2005; Proks et al., 1999; Tammaro et al., 2005; Tucker et al., 1998), consistent with a role of these residues in ATP gating. Notably, sequence comparison reveals that a key difference between Kir6.2 and other Kir channels is G334, which in other Kir channels is occupied by larger amino acids. Substitution of glycine at this position by a larger amino acids such as histidine seen in Kir2 or Kir3 channels would create steric hindrance to prevent ATP binding. This may explain, at least in part, why Kir6.2 is the only Kir channel sensitive to ATP regulation.

### Structural interactions around the ATP binding site and their relationship to the PIP_2_ binding site

A number of residues within the vicinity of the ATP-binding site such as E179, R201, and R301 have previously been shown to reduce ATP sensitivity and proposed to be involved in ATP binding (Haider et al., 2005). However, from the structure it is clear these residues contribute indirectly (Fig.5). E179 and R301 were both proposed to interact with the adenine ring (Haider et al., 2005). In our structure, neither residue forms direct interactions with ATP. E179 appears to interact with R54 from the adjacent Kir6.2 and may be part of the network that stabilizes the interaction between R50 and ATP (Fig.5C). R301 is found to interact with Q299 in the same β-strand that is part of a β-sheet in the Kir6.2 CTD (Fig. 5A; see also Fig.2-figure supplement 1). Interestingly, R301 is one of the most highly mutated residues in congenital hyperinsulinism; mutation of R301, in addition to mildly reducing ATP sensitivity, results in rapid decay of channel activity that can be reversed by increasing PIP_2_ concentrations in the membrane (Lin et al., 2008). Based on our structure, it is likely that mutation of this residue disrupts structural integrity of the Kir6.2 CTD necessary for stable channel interaction with PIP_2_ and ATP. By contrast, R201 is one of the most highly mutated residues in neonatal diabetes (Ashcroft, 2005). It has been proposed that R201 coordinates the α-phosphate of ATP (Haider et al., 2005). However, in the structure R201 is found on the β-strand directly below that of I182 and K185, and is too distant to directly interact with ATP. Instead, R201 is sandwiched between the benzene rings of F333 and F315, forming a dual cation-π interaction that likely stabilizes the ATP-binding site (Fig. 5B). Mutation of R201 would therefore destabilize this interaction to indirectly reduce ATP inhibition.

**Fig. 5.**
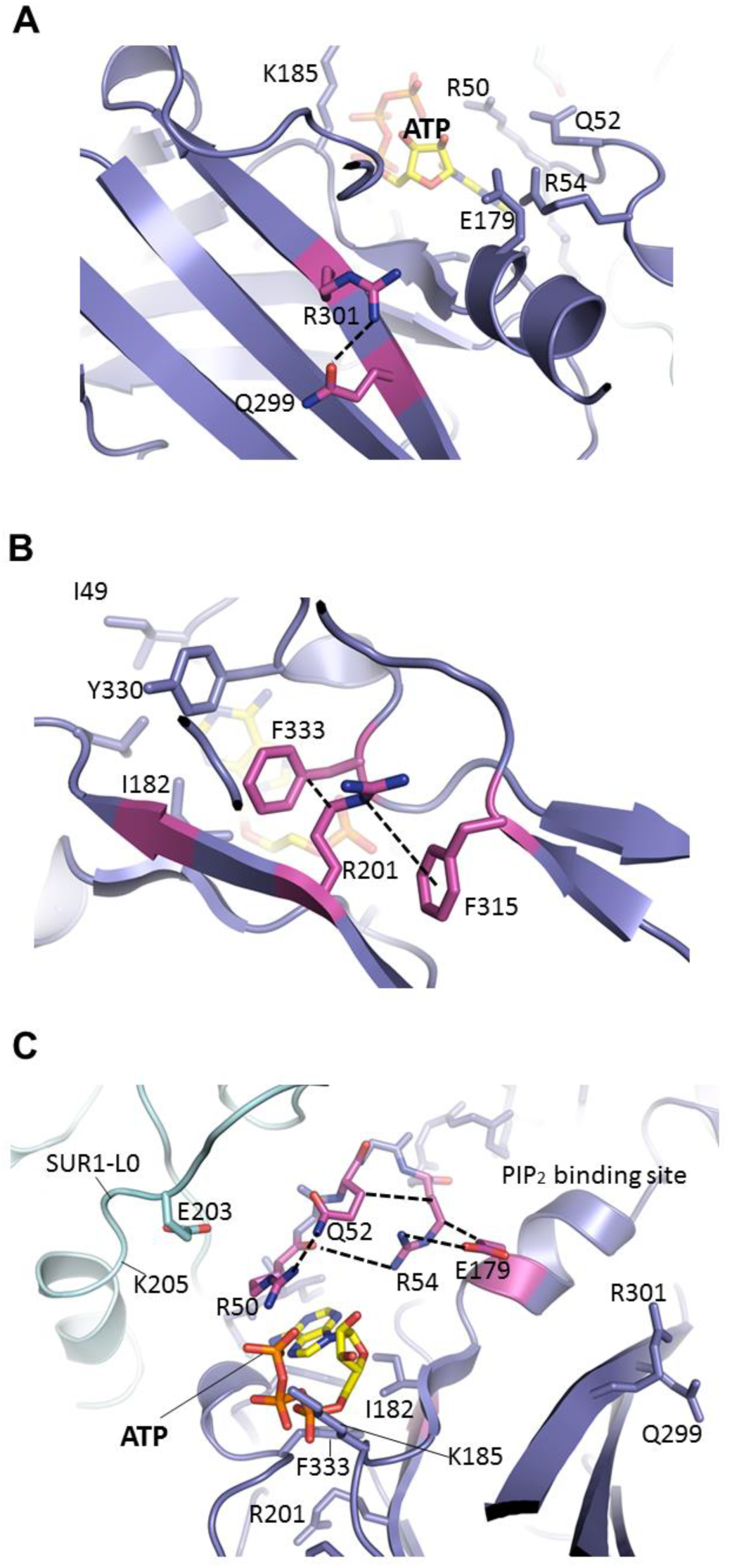
Important molecular interactions surrounding the ATP binding site. (**A**) Electrostatic interaction between R301 and Q299, viewed from the interior of Kir6.2 and looking out toward the cytoplasm. These residues are found on an internal β-sheet 12Å from ATP (Cα of R301 to ribose of ATP) (**B**) Dual interaction between R201 and F315, likely via cation-π, and hydrophobic stacking between aliphatic portion of R201 side chain and F333, again viewed from the Kir6.2 interior. These residues are found directly below ATP (~9Å from Cα of R201 to ribose of ATP).

Another interesting residue is Q52. The PNDM mutation Q52R causes extremely high *P_o_* and very low ATP sensitivity (Lin et al., 2006; Proks et al., 2004). In the structure, Q52 interacts with R50 which coordinates the γ-phosphate of ATP, but is also interacting with R54, which orients the R54 side chain towards the ATP site and away from the PIP_2_ site nearby (Fig.5C). In the PIP_2_ bound Kir2.2 structure, the Kir6.2 R54 equivalent arginine residue interacts with the tether helix near PIP_2_ binding residues (Hansen et al., 2011), suggesting that R54 may be important for Kir6.2-PIP_2_ interactions. Interestingly, mutation of R50 or R54 to an alanine has been reported to reduce sensitivity to both ATP and PIP_2_ (Cukras et al., 2002). From our structure, it is easy to envision how mutation of any of these residues can disrupt the interaction network to affect gating by either ligand.

It is also important to note that in our structure, Q52 is in close proximity to E203 in the L0 of SUR1 immediately following TMD0. We have previously shown that engineered interactions between Kir6.2 residue 52 and SUR1 residue 203 via a Kir6.2-Q52E and SUR1-E203K ion pair increases channel sensitivity to ATP by nearly two orders of magnitude, and that crosslinking of the two residues via a Kir6.2-Q52C and SUR1-E203C mutant pair induces spontaneous channel closure in the absence of ATP (Pratt et al., 2012). In addition, our previous studies have shown that ATP binding involves residues from not only the N-terminus of Kir6.2 such as R50 but also residues in SUR1-L0 such as K205 (Martin et al., 2017; Pratt et al., 2012). Together these studies lead us to propose that the inhibitory effect of ATP is partially due to stabilizing the interaction between this N-terminal region of Kir6.2 and L0 of SUR1 (see Discussion).

### The GBC binding site

Sulfonylureas stimulate insulin secretion by inhibiting pancreatic K_ATP_ channels. GBC, also known as glyburide, is a second generation sulfonylurea that contains both a sulfonylurea moiety and a benzamido moiety, and binds K_ATP_ channels with nM affinity (Gribble and Reimann, 2003). Despite intense investigation the GBC binding site has remained elusive. Early studies using chimeras of SUR1 and SUR2A, which is known to have lower sensitivity to GBC than SUR1, suggest the involvement of TMs 14-16; in particular mutating S1238 in SUR1 to Y (note in some papers, this is numbered as S1237) as seen in SUR2A compromised GBC binding and block (Ashfield et al., 1999; Winkler et al., 2007). Subsequent studies using ^125^I-azido-GBC photolabeling implicated involvement of L0 of SUR1; specifically, two mutations Y230A and W232A in L0 severely compromised photolabeling of SUR1 (Vila-Carriles et al., 2007). These studies led to a model in which S1238 and Y230/W232 constitute two ends of a bipartite binding pocket, each recognizing opposite ends of GBC (Winkler et al., 2007); however, whether one or both contribute directly to GBC binding is unknown.

In the current reconstruction we find well defined, non-protein density within the TMDs of SUR1, with a size and shape which closely matches that of a GBC molecule (Fig.6, Fig.6-figure supplement 1A). One end of the density is in direct contact with S1238 and resembles the cyclohexyl moiety long presumed to constitute the “A” site that is abolished by the S1238Y mutation (Ashfield et al., 1999; Bryan et al., 2004). We used this to guide the initial docking of GBC, which could then be readily refined into the density together with SUR1.

**Fig. 6.**
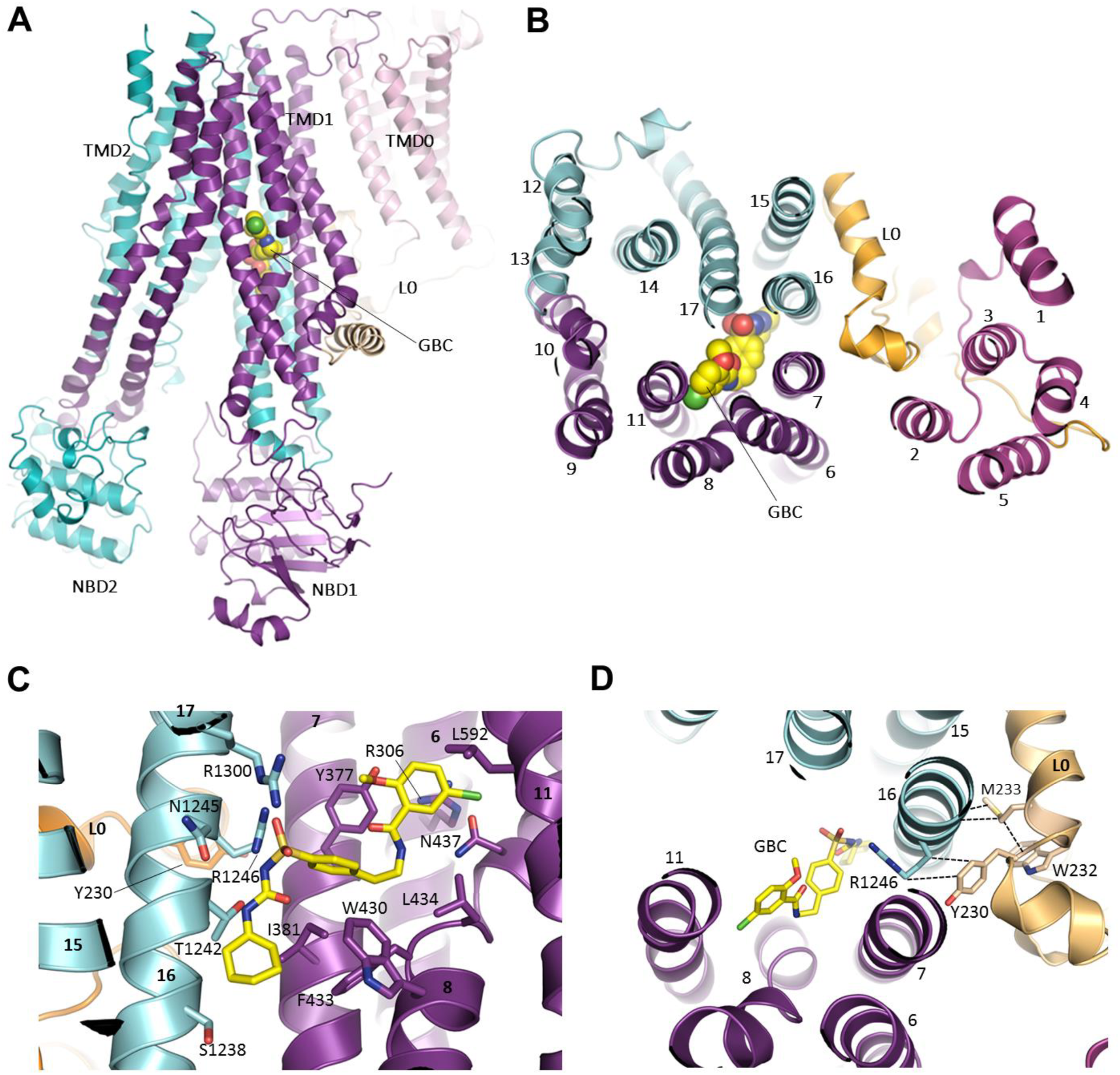
The GBC binding site. (**A**) Ribbon diagram of SUR1 showing location of GBC, which is primarily coordinated by the inner helices of TMD1 (purple) and TMD2 (cyan). (**B**) Slice view of model in (**A**) viewed from the extracellular side. Note juxtaposition of L0 to helices in ABC core directly interacting with GBC. (**C**) Close-up of GBC binding pocket, showing all residues which immediately line the pocket and seem to form direct contact with GBC; a subset of these residues were mutated to test their role in GBC binding (Fig. 7). (**D**) Magnified view in (**B**), highlighting indirect roles of Y230 and W232 (L0) in GBC binding. These both likely stabilize interaction between residues on helix 16 of TMD2 and GBC, at the same time anchoring this helix of L0 to the ABC core structure.

The binding pocket is contoured to precisely accommodate GBC, and the combination of polar and hydrophobic residues help explain the sub-nM affinity of SUR1 for this sulfonylurea (Fig. 6C, Fig.6-figure supplement 1B, C). A primary anchor is composed of two arginine residues, R1246 and R1300, which coordinate each oxygen of the sulfonyl group. Each nitrogen of the urea moiety is coordinated by T1242 and N1245, and the adjacent benzene and cyclohexyl groups (adjacent to the sulfonyl and urea groups, respectively) are stabilized by a series of hydrophobic interactions contributed by TM helices from both TMD1 (TM6, 7, 8) and TMD2 (TM16). As a second-generation sulfonylurea, GBC contains another lipophilic group adjacent to an amide linker, which is lacking in first-generation compounds like tolbutamide. This group, a 1-chloro-4-methoxy-benzene, is encircled by a ring of hydrophilic and hydrophobic side chains. In particular, the Cl appears to hydrogen bond with the amino group of N437, while the methoxy is H-bonded to the hydroxyl group of Y377. Y377 also seems to contribute a π-π stacking interaction with the benzene ring. In the structure, the previously proposed sulfonylurea binding residue S1238 juxtaposes the cyclohexyl group of GBC, with only ~3Å separation between the Cβ of S1238 and the 6-carbon ring of GBC. Mutation of this residue to a tyrosine may alter interaction with GBC to compromise high affinity binding of GBC.

In order to validate the proposed binding site, we mutated a subset of key GBC-binding residues listed above to alanine and tested their response to 100 nM and 1µM GBC with Rb^+^ efflux experiments, which measure channel activity and response to GBC in intact cells. All six mutations, R306A, Y377A, N437A, T1242A, R1246A, and R1300A trafficked normally and responded as WT to metabolic inhibition (Fig.7A, B). Strikingly, all six mutations showed significantly reduced or complete absence of inhibition at 100 nM, and five mutations (R306A, Y377, N437A, T1242, and R1246A) showed significantly reduced sensitivity compared to WT even at 1µM GBC (Fig. 7C, D). The four most GBC-insensitive mutants, R306A, Y377A, N437A, and T1242A were further analyzed by inside-out patch-clamp recording. Although these mutants were still sensitive to GBC inhibition, the extent of inhibition at steady-state was less compared to WT channels at 10nM and 100nM (Fig.7, figure supplement 1). Also worth noting, while inhibition of WT channels was nearly irreversible, inhibition of mutants was more reversible, consistent with the mutants having reduced affinity for GBC (Fig.7-figure supplement 1). Together these results provide strong functional evidence for the GBC binding pocket defined in our structure.

**Fig. 7.**
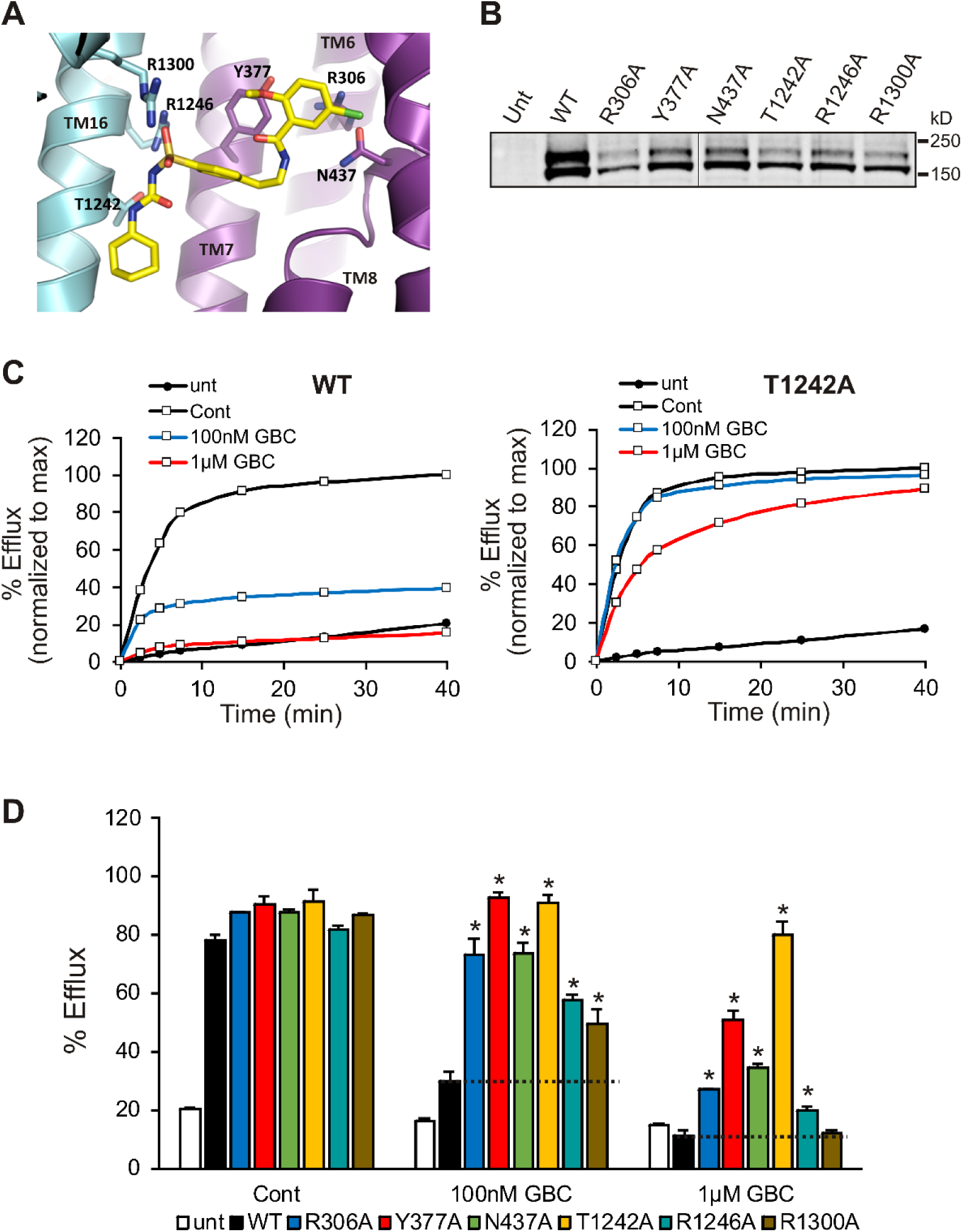
Functional testing of GBC binding residues. **(A)** Residues in SUR1 selected to be mutated to alanine. **(B)** Western blot of WT and mutant SUR1 co-expressed with Kir6.2 in COS cells. Two SUR1 bands corresponding to the core-glycosylated immature protein (lower band) and the complex-glycosylated mature protein (upper band) are detected. The vertical line in the middle of the blot separates two parts of the same blot. **(C)** Representative efflux profiles of WT channels and T1242A mutant channels in cells pretreated with metabolic inhibitors for 30 min in the presence of 0.1% DMSO (cont), 100nM GBC, or 1µM GBC. Untransfected cells (unt) served as a control. Efflux was normalized to the maximal value observed at 40 min for direct comparison. **(D)** Quantification of percent efflux of all mutants compared to WT. Each bar represents the mean±s.e.m. of 3-4 experiments. * *p*<0.05 by one-way ANOVA with Newman-Keuls *post hoc* test.

### The role of SUR1-Y230 and W232 in GBC interaction

In the previously proposed bipartite binding model for GBC, the pocket was formed from two overlapping regions: at one end was S1238 of TMD2, and at the other was L0 involving residues Y230 and W232, part of the “lasso motif” observed in MRP1 and CFTR. Mutation of Y230 to an alanine has also been shown to reduce the ability of GBC to inhibit channel activity (Devaraneni et al., 2015; Yan et al., 2006). In our current structure, Y230 is too distant to interact directly with GBC. However, the binding pocket is close to the L0-TMD interface, where the L0 amphipathic helix forms a series of mostly hydrophobic interactions with transmembrane helices from TMD1/2 that line the GBC binding pocket. Here, we find that Y230 stacks closely against the aliphatic portion of the R1246 side chain, which in turn coordinates an oxygen of the sulfonyl group of GBC (Fig.6D). W232 appears to form a strong interaction with M233, which interacts directly with two alanines, A1243 and A1244, on the opposite side of TM16 where two GBC interacting residues T1242 and N1245 are located (Fig.6D). These observations indicate an important but clearly indirect role for Y230 and W232 in GBC binding.

## Discussion

The structure presented in this study is the first to reveal in detail the ATP and GBC binding sites in the SUR1/Kir6.2 K_ATP_ channel complex. The clear density for ATP and GBC as well as all residues involved in binding of both ligands in the current EM map allowed us to present a detailed atomic interpretation of ATP and GBC binding to the channel. Importantly, the binding pockets we identified are supported by strong functional data. In addition, the structure uncovers many molecular interactions that indirectly impact ATP and GBC gating, and those that underlie SUR1-Kir6.2 interactions. The structural information gained offers key insights into possible mechanisms of how the two ligands both inhibit K_ATP_ channels to stimulate insulin secretion.

### The ATP binding site and mechanism of ATP inhibition

The Kir6.2 interfacial ATP binding site model was first proposed by Antcliff et al. (Antcliff et al., 2005) based on ligand docking, homology modeling of Kir channel crystal structures, and structure-function mutagenesis data. Although some interactions in the original model between Kir6.2 residues and ATP are observed in our structure, many others require new interpretations. First, the ATP molecule adopts a conformation with the γ-phosphate bent towards the adenine ring (Fig.4), which is reminiscent of that observed in P2X receptors, an ATP activated ion channel (Hattori and Gouaux, 2012). Second, our structure suggests that E179, R201, and R301 rather than contributing directly to ATP binding are critical for interactions with other residues that support the ATP binding residues or general structural integrity of the Kir6.2 CTD for stable channel interaction with PIP_2_ (Fig.5). Elucidation of the structural role of R201 and R301 helps us to understand the mechanisms by which mutation of these residues cause insulin secretion disease. Finally, the structure provides details on the intricate relationship between ATP binding residues and those which have been implicated in PIP_2_ binding or gating, shedding light on how the channel senses ATP and PIP_2_ (Fig.5C).

We propose that ATP, by binding to a pocket created by the N-terminus and CTD from two adjacent Kir6.2 subunits with contributions from L0 of SUR1, acts like a stopper that prevents movement of the Kir6.2 N-terminus relative to L0 that is necessary to open the channel (Fig. 8A). This model is consistent with our previous study showing that crosslinking of SUR1-L0 with the N-terminus of Kir6.2 near the ATP binding site locks the channel closed even without ATP (Pratt et al., 2012). One interesting question is whether the interaction network we observed in the present ATP-bound structure undergoes remodeling in PIP_2_-bound open state. An open state structure of the K_ATP_ channel bound to PIP_2_ will be needed to understand the full extent of conformational change associated with ATP and PIP_2_ gating.

**Fig. 8.**
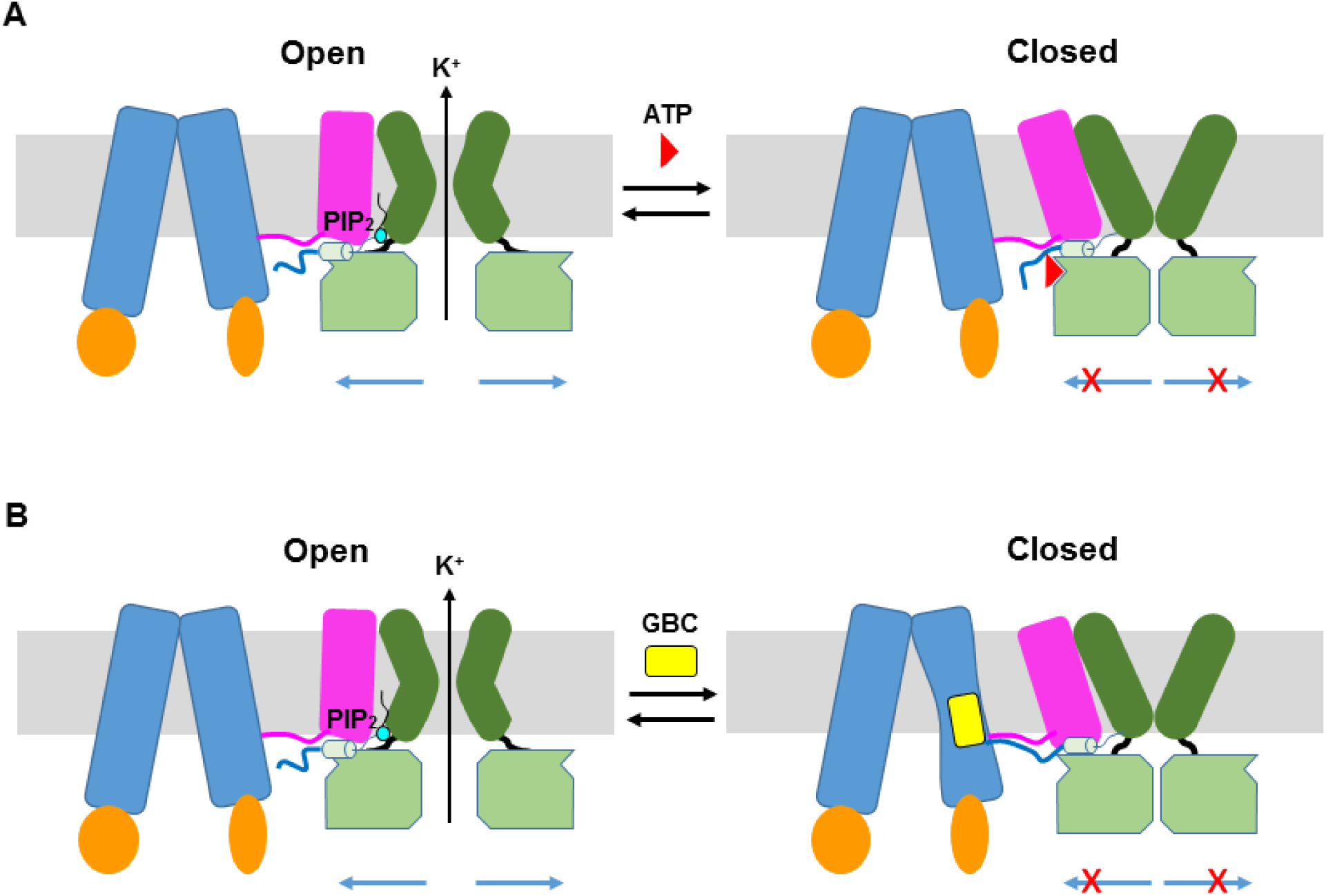
ATP and GBC gating models. **(A)** Cartoon model illustrating that ATP binds to a pocket formed by the N-terminus and CTD of Kir6.2 (from two adjacent subunits), with contribution form L0 of SUR1. This prevents movement of the Kir6.2 N-terminus that is necessary to open PIP_2_-bound channels. (**B**) Hypothetical model of GBC gating. GBC binds to the TM bundle juxtaposing L0, which stabilizes the SUR1-ABC core in an inward-facing conformation. GBC also stabilizes interactions of the distal N-terminus of Kir6.2 with SUR1 thus restricting movement of the Kir6.2 N-terminus to inhibit channel activity. In both A and B, Kir6.2 transmembrane helices: dark green; Kir6.2 cytoplasmic domain: green; Kir6.2 slide helix and N-terminus from adjacent subunit: light green cylinder and thick blue line, respectively; SUR1-TMD0/L0: magenta; SUR1-TMD1/2: blue; SUR1-NBDs: orange; GBC: yellow; ATP: red; PIP_2_: cerulean. Note the different states shown are not meant to reflect the actual conformational transitions. Also, PIP_2_ was not included in the closed states; however, it remains to be determined whether ATP precludes PIP_2_ binding.

### Mechanistic insights of GBC binding and inhibition

The binding site of the high affinity sulfonylurea GBC has been studied by many groups. These studies have implicated the involvement of transmembrane helices in the SUR1-ABC core, L0, and the N-terminus of Kir6.2 (Ashfield et al., 1999; Bryan et al., 2004; Vila-Carriles et al., 2007). Yet, the precise binding pocket for this commonly used anti-diabetic drug has remained unresolved. In a recent 5-6Å resolution reconstruction of a pancreatic K_ATP_ channel complex imaged in the presence of GBC, Li et al. (Li et al., 2017) assigned extra density around Y230 and W232 of SUR1 as GBC, thus placing the sulfonylurea binding site within an amphipathic helix of L0. However, at low resolution modeling of L0 is challenging, as this is an inherently lower resolution part of the structure; moreover, because Y230 and W232 are found at the interface of the inner leaflet of the bilayer, bound lipids could easily be mistaken for ligand density. Contrary to the study by Li et al., we were able to clearly assign the GBC density in the TM bundle connected to the NBD1, with residues from TM6, 7, 8, 11 in TMD1 and TM16 and 17 from TMD2 contributing to GBC interactions. Importantly, our model is supported by functional data using both ^86^Rb^+^ efflux assays and electrophysiological recordings. Moreover, our structure clarifies how Y230 and W232, which have previously been proposed to contribute to GBC binding based on indirect biochemical or functional assays (Vila-Carriles et al., 2007), can affect GBC binding or gating indirectly by supporting residues that are directly engaged in GBC binding.

The distal N-terminal 30 amino acids of Kir6.2 have been shown to be important for the binding or effect of GBC in a number of studies (Devaraneni et al., 2015; Koster et al., 1999; Kuhner et al., 2012; Reimann et al., 1999; Vila-Carriles et al., 2007). In our map, there is a lack of strong density N-terminal to position 32 of Kir6.2, suggesting this region is flexible. However, it is worth noting that our previous study using engineered unnatural amino acid Azido-*p-*phenylalanine placed at the distal N-terminus of Kir6.2 (amino acid position 12 or 18) has demonstrated that GBC increased crosslinking of Kir6.2 to SUR1 (Devaraneni et al., 2015). Accordingly, we propose that GBC binding to the TMD bundle next to the L0 amphipathic helix stabilizes SUR1 in an inward-facing conformation, and possibly also stabilizes the interactions between the N-terminus of Kir6.2 and SUR1, as we have demonstrated previously, to prevent the movement of Kir6.2 N-terminus that is needed to open the gate (Fig.8B).

The residues which play a specific and critical role in GBC binding are also very likely important for binding of other sulfonylureas. While we only tested GBC, we predict that R1246 and R1300, which coordinate the sulfonyl group, and T1242 and N1245, which coordinate the urea group, will also be critical for binding of other sulfonylureas such as tolbutamide (Gribble and Reimann, 2003). In addition to sulfonylureas, glinides such as rapaglinide and nateglinide which lack the sulfonylurea moiety (Gribble and Reimann, 2003), and a structurally unrelated compound carbamazepine (Chen et al., 2013b; Devaraneni et al., 2015) are also known to inhibit K_ATP_ channels. Elucidating the role of the various GBC binding residues in channel interactions with the different channel inhibitors will be important for understanding channel inhibition mechanisms and for rational design of new drugs with desired properties.

### Conservation of GBC binding residues in other SUR proteins and ABCC transporters

Multiple sequence alignment of 15 SUR1 orthologs from diverse genera shows relatively high sequence identity throughout the sequence relative to human; from 95% (hamster) to 75% (seahorse). Interestingly, the segments of the helices from TMD1 and TMD2 which comprise the GBC binding site show exceptionally high conservation, with every one of the 12 residues which most closely line the GBC pocket absolutely conserved in 15 out of the 15 sequences. The high degree of conservation suggests the importance of the interface formed by these transmembrane helices. It would be important to determine in the future whether this interface is involved in conformational switch of the SUR1 ABC core and its communication with Kir6.2.

Interestingly, SUR2 (*ABCC9*), the closest homolog of SUR1 (67% sequence identity), while also shows high conservation of the GBC binding residues (10/12), differs in two positions that correspond to S1238 and T1242 (Y and S respectively in SUR2). SUR2 assembles with Kir6.1 or Kir6.2 to form K_ATP_ channel subtypes found in the heart, skeletal muscle, and vascular smooth muscle. These channels are known to have lower sensitivity to GBC inhibition than channels formed by SUR1 and Kir6.2 (Inagaki et al., 1996). Variations at these two key GBC binding residues likely explains their different pharmacological sensitivity to GBC (Ashfield et al., 1999; Inagaki et al., 1996).

In addition to targeting K_ATP_ channels, GBC has been shown to inhibit other ABC transporters within the C subfamily (ABCC), including MRP1 and CFTR, albeit at lower affinity (~30µM for both CTFR and MRP1) (Payen et al., 2001; Schultz et al., 1996). Of all the residues within the GBC binding pocket, only two appear to be highly conserved across different members of the ABCC subfamily: R1246 and R1300. In fact, R1246 is strictly conserved within 11 of 12 ABCC homologs (Gln in ABCC10), and R1300 in 10 of 12 (Asn in CFTR and Ser in ABCC10). In the cryo-EM structure of MRP1 bound to substrate LTC-4 (Johnson and Chen, 2017), R1196 (equivalent to R1246 in SUR1) forms a salt bridge with a carboxylic acid group of LTC-4 and is also in the same rotameric conformation as R1246 in SUR1. Further, F221 (equivalent to Y230 in SUR1) also seems to form the equivalent hydrophobic stacking interaction with R1196 as Y230 does with R1246 in SUR1; this phenomenon is also observed in the human CFTR structure (F17 and R1097) (Liu et al., 2017). Such structural conservation likely explains the GBC sensitivity in other ABCC homologues, and suggests a critical role for this pair of residues in the function and/or structure of ABCC proteins.

In summary, we presented a K_ATP_ channel structure with improved resolution that allowed us to definitively identify the ATP and GBC binding sites. The novel insight gained from this structure significantly advances our understanding of how these two ligands interact with the channel to exert an inhibitory effect. As inhibition of K_ATP_ channels by sulfonylureas remains an important therapeutic intervention to control type 2 diabetes and neonatal diabetes, and there is a need for drugs that specifically target a K_ATP_ channel subtype, our study offers a starting point for future structure-guided drug development to mitigate diseases caused by K_ATP_ channel dysfunction.

## Materials and methods

### Protein expression and purification

K_ATP_ channels were expressed and purified as described previously (Martin et al., 2017). Briefly, the genes encoding pancreatic K_ATP_ channel subunits, which comprise a hamster SUR1 and a rat Kir6.2 (94.5% and 96.2% sequence identity to human, respectively), were packaged into recombinant adenoviruses (Lin et al., 2005; Pratt et al., 2009); Both are WT sequences, except for a FLAG tag (DYKDDDDK) that had been engineered into the N-terminus of SUR1 for affinity purification. INS-1 cells clone 832/13 (Hohmeier et al., 2000), a rat insulinoma cell line, were infected with the adenoviral constructs in 15 cm tissue culture plates. Protein was expressed in the presence of 1mM Na butyrate and 5 μM GBC to aid expression of the channel complex. 40-48 hours post-infection, cells were harvested by scraping and cell pellets were frozen in liquid nitrogen and stored at −80°C until purification.

For purification, cells were resuspended in hypotonic buffer (15mM KCl, 10mM HEPES, 0.25 mM DTT, pH 7.5) and lysed by Dounce homogenization. The total membrane fraction was prepared, and membranes were resuspended in buffer A (0.2M NaCl, 0.1M KCl, 0.05M HEPES, 0.25mM DTT, 4% sucrose, 1mM ATP, 1uM GBC, pH 7.5) and solubilized with 0.5% Digitonin. The soluble fraction was incubated with anti-FLAG M2 affinity agarose for 4 hours and eluted with buffer A (without sugar) containing 0.25 mg/mL FLAG peptide. Purified channels were concentrated to ~1-1.5 mg/mL and used immediately for cryo grid preparation.

### Sample preparation and data acquisition for cryo-EM analysis

In our previous data sets, most micrographs were of ice that was either too thin, which tended to exclude channel complex from the hole and induce highly preferred orientation, or of ice that was too thick, which gave good particle distribution and good angular coverage, but had lower contrast. Thus the current data set was the result of efforts to optimize ice thickness in order to retain high contrast and particle distribution. This was achieved through varying blotting time and also through extensive screening of the grid in order to find optimal regions. Two grids were imaged from the same purification and were prepared as follows: 3 µL of purified K_ATP_ channel complex was loaded onto UltrAufoil gold grids which had been glow-discharged for 60 seconds at 15 mA with a Pelco EasyGlow ®. The sample was blotted for 2s (blot force −4; 100% humidity) and cryo-plunged into liquid ethane cooled by liquid nitrogen using a Vitrobot Mark III (FEI).

Single-particle cryo-EM data was collected on a Titan Krios 300 kV cryo-electron microscope (FEI) in the Multi-Scale Microscopy Core at Oregon Health & Science University, assisted by the automated acquisition program SerialEM. Images were recorded on the Gatan K2 Summit direct electron detector in super-resolution mode, post-GIF (20eV window), at the nominal magnification 81,000x (calibrated image pixel-size of 1.720Å; super-resolution pixel size 0.86Å); defocus was varied between −1.4 and −3.0 µm across the dataset (Table 1). The dose rate was kept around 2.7 e^-^/Å^2^/sec, with a frame rate of 4 frames/sec, and 60 frames in each movie, which gave a total dose of approximately 40 e^-^/Å^2^. In total, 2180 movies were recorded.

### Image processing

The raw frame stacks were gain-normalized and then aligned and dose-compensated using Motioncor2 (Zheng et al., 2017) with patch-based alignment (5x5). CTF parameters were estimated from the aligned frame sums using CTFFIND4 (Rohou and Grigorieff, 2015). Particles were picked automatically using DoGPicker (Voss et al., 2009) with a broad threshold range in order to reduce bias. Subsequently, each image was analyzed manually in order to recover any particles missed by automatic picking and remove obviously bad micrographs from the data set. This resulted in ~250,000 raw particles as input for subsequent 2D classification using RELION-2 (Kimanius et al., 2016). After four rounds of 2D classification, ~160,000 particles remained in the data set, in which only classes displaying fully assembled complexes and high signal/noise were selected. These 160K particles were re-extracted at 1.72 Å/pix and were used as input for 3D classification in RELION-2.

Extensive 3D classification was performed in order to sample the heterogeneity within the data. Symmetry was not imposed at this step in order to select only true four-fold classes. Up to 4 consecutive rounds of classification were performed, specifying 4 or 5 classes per round. Individual classes and combinations of classes were refined independently and lead to very similar structures. The two best classes from round 2 were combined (~63,000 particles), and then particles were re-extracted from super-resolution micrographs with a box size of 600 pixels. A soft mask encompassing the entire complex was used during refinement in RELION, with C4 symmetry imposed, which resulted in a 3.8Å reconstruction using the gold-standard FSC cutoff. The map was B-factor corrected and filtered using RELION-2 Postprocessing procedure, with the same mask used for refinement. Local resolution was calculated on the unfiltered map with the Bsoft package, which showed the resolution was highest in the Kir.6.2/TMD0 core, as well as the SUR1 helices surrounding the GBC-binding pocket (between 3.3-3.7), and lowest in the NBDs and most external helices of TMD1/TMD2 of SUR1 (Fig.1-figure supplement 2).

### Model building

In our previous reconstruction, many side-chains were left out of the final model as there was not sufficient density to support their placement (Martin et al., 2017). In the current reconstruction, there is good density for nearly every side chain of Kir6.2, TMD0, and the inner helices of the ABC core structure of SUR1. Thus using our previous structure as the starting template, we rebuilt nearly all of the structure with RosettaCM (Song et al., 2013), using the density as an additional constraint. This region included Kir6.2, TMD0/L0, and TMD1 and TMD2. The lowest energy models were very similar to one another, thus the lowest energy model was selected for each region. The resulting model was then minimized once in CNS (Brunger et al., 1998), substituting in the RSRef real-space target function (Chapman et al., 2013), adding (φ,ψ) backbone torsion angle restraints, and imposing non-crystallographic symmetry (NCS) constraints. In the density map, NBD1 and NBD2 showed signs of disorder, so our previously deposited NBD models were left as polyalanine chains and only refined as rigid bodies with RSRef. The distal N- and C-termini of Kir6.2, as well as the linker between NBD1 and TMD2 in SUR1 were not observed in the density map, and thus were left out the model. The final model contains residues 32-352 for Kir6.2, and residues 6-615 (TMD0/L0 + TMD1), 678-744 and 770-928 (NBD1), 1000-1044 and 1061-1319 (TMD2), and 1343-1577 (NBD2) for SUR1. All structure figures were produced with UCSF Chimera (Pettersen et al., 2004) and PyMol (http://www.pymol.org). Pore radius calculations were performed with HOLE (Smart et al., 1996).

### Sequence alignments

Multiple sequence alignment was performed with the T-Coffee server (Notredame et al., 2000). Output was saved in Clustal Aln format, and then imported and visualized in UCSF Chimera.

### Functional studies of GBC binding mutants

Point mutations were introduced into hamster SUR1 cDNA in pECE using the QuikChange site-directed mutagenesis kit (Stratagene). Mutations were confirmed by DNA sequencing. Mutant SUR1 cDNAs and rat Kir6.2 in pcDNA1 were co-transfected into COS cells using FuGENE®6, as described previously (Devaraneni et al., 2015) and used for Western blotting, ^86^Rb^+^ efflux assays, and electrophysiology as described below.

For Western blotting, cells were lysed in 20 mM HEPES, pH 7.0/5 mM EDTA/150 mM NaCl/1% Nonidet P-40 with CompleteTR protease inhibitors (Roche) 48-72 hours post-transfection. Proteins in cell lysates were separated by SDS/PAGE (8%), transferred to nitrocellulose membrane, probed with rabbit anti-SUR1 antibodies against a C-terminal peptide of SUR1 (KDSVFASFVRADK), followed by HRP-conjugated anti-rabbit secondary antibodies (Amersham Pharmacia), and visualized by chemiluminescence (Super Signal West Femto; Pierce) with FluorChem E (ProteinSimple).

For ^86^Rb^+^ efflux assays, cells were plated and transfected in 12-well plates. Twenty-four to thirty-six hours post-transfection, cells were incubated overnight in medium containing ^86^RbCl (0.1 μCi/ml). The next day, cells were washed in Krebs-Ringer solution twice and incubated with metabolic inhibitors (2.5μg/ml oligomycin and 1mM 2-deoxy-D-glucose) in Krebs-Ringer solution for 30 min in the presence of ^86^Rb^+^. Following two quick washes in Krebs-Ringer solutions containing metabolic inhibitors and 0.1% DMSO (vehicle control), 100nM GBC, or 1µM GBC, 0.5 ml of the same solution was added to each well. At the end of 2.5 minutes, efflux solution was collected for scintillation counting and new solution was added. The steps were repeated for 5, 7.5, 15, 25, and 40 min cumulative time points. After the 40 min time point efflux solution was collected, cells were lysed in Krebs-Ringer containing 1% SDS. ^86^Rb^+^ in the solution and the cell lysate was counted. The percentage efflux was calculated as the radioactivity in the efflux solution divided by the total activity from the solution and cell lysate, as described previously (Chen et al., 2013a; Yan et al., 2007). Note we used higher concentrations of GBC for these experiments than the electrophysiology experiments described below as in the latter the channels were exposed directly in isolated membrane patches to GBC, thus requiring lower concentrations. Experiments were repeated three-four times and for each experiment, untransfected cells were included as a negative control.

For electrophysiology experiments, cells co-transfected with SUR1 and Kir6.2 along with the cDNA for the green fluorescent protein GFP (to facilitate identification of transfected cells) were plated onto glass coverslips twenty-four hours after transfection and recordings made in the following two days. All experiments were performed at room temperature as previously described (Devaraneni et al., 2015). Micropipettes were pulled from non-heparinized Kimble glass (Fisher Scientific) on a horizontal puller (Sutter Instrument, Co., Novato, CA, USA). Electrode resistance was typically 1-2 MΩ when filled with K-INT solution containing 140 mM KCl, 10 mM K-HEPES, 1 mM K-EGTA, pH 7.3. ATP was added as the potassium salt. Inside-out patches of cells bathed in K-INT were voltage-clamped with an Axopatch 1D amplifier (Axon Inc., Foster City, CA). ATP (as the potassium salt) or GBC at 10nM or 100nM were added to K-INT as specified in the figure legend. All currents were measured at a membrane potential of −50 mV (pipette voltage = +50 mV). Data were analyzed using pCLAMP10 software (Axon Instrument). Off-line analysis was performed using Microsoft Excel programs. Data were presented as mean±standard error of the mean (s.e.m).

## Acknowledgements

The INS-1 cell clone 832/13 was kindly provided by Dr. Christopher Newgard. We thank Emily Rex and Zhongying Yang for technical assistance, and Dr. Matt Whorton, Dr. Dale Fortin, and Veronica Cochrane for critical readings of the manuscript. We also thank the staff at the Multiscale Microscopy Core (MMC) of Oregon Health & Science University (OHSU), the OHSU-FEI living lab and Intel for technical support. This work was supported by the National Institutes of Health grants R01DK066485 (to S.-L. S.) and F31DK105800 (to G.M.M.).

## Author contributions

GMM, Data curation, Formal analysis, Funding acquisition, Validation, Investigation, Visualization, Methodology, Writing—original draft, Writing—review and editing; CY, Data curation, Software, Formal analysis, Validation, Visualization, Writing—review and editing; BK, Data curation, Investigation, Writing—review and editing; FD, Software, Validation, Writing— review and editing; SLS, Data curation, Formal analysis, Funding acquisition, Validation, Investigation, Visualization, Methodology, Writing—original draft, Writing—review and editing.

---

**Fig. 1-figure supplement 1.**
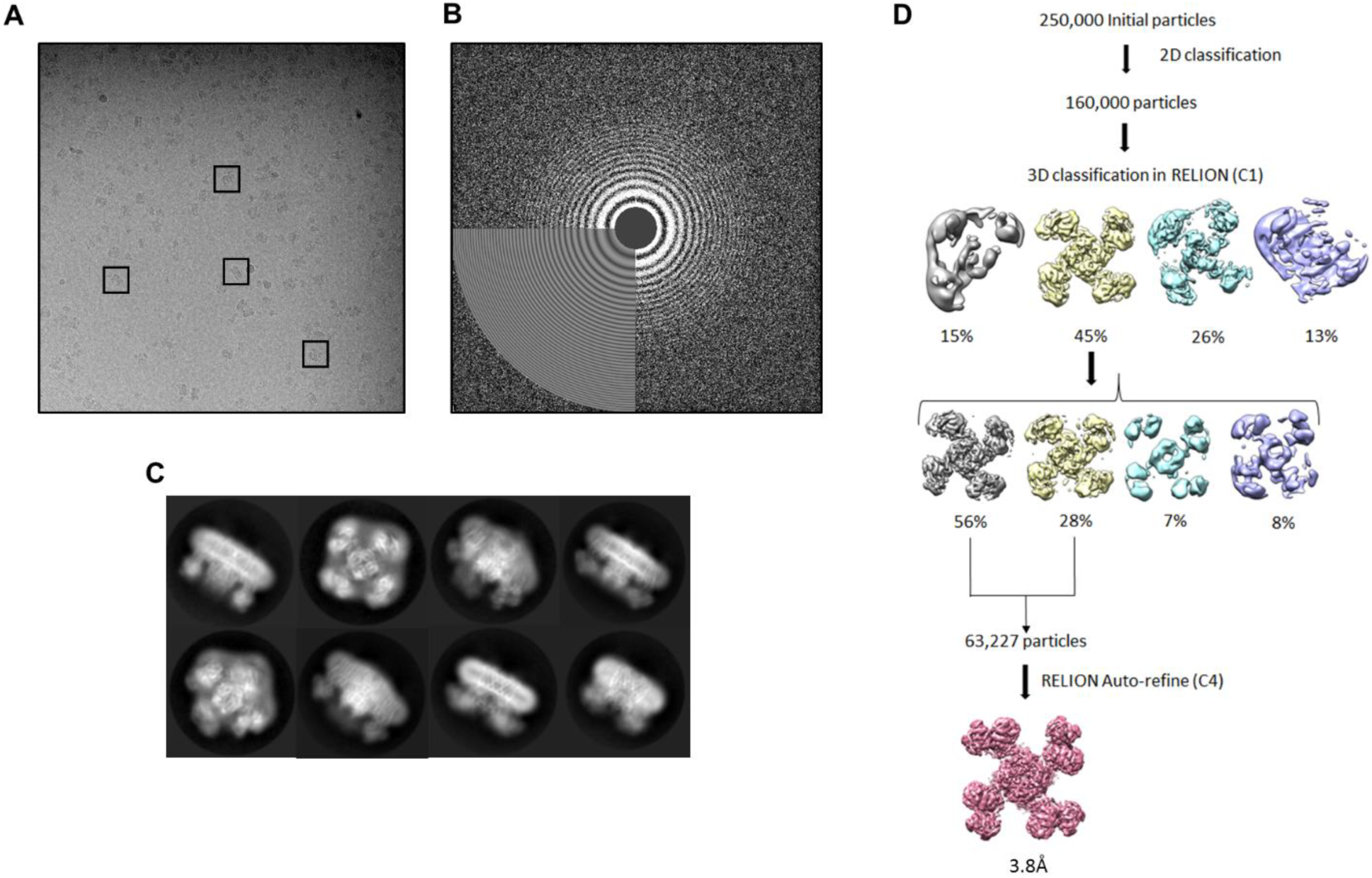
Data collection and image processing workflow. **(A)** Representative micrograph at 81,000x (1.72 Å/pixel; 0.86 Å/pixel super-resolution) after alignment with Motioncor2. A few K_ATP_ channel complexes of various orientation have been outlined. **(B)** Power spectrum calculated with Ctffind4, with information extending out to 3.6Å. **(C)** Select 2D classes from final round of classification. **(D)** Overview of data processing workflow. Particle picking was performed automatically with DogPicker as well as with manual inspection. All other image processing steps were performed in RELION-2.

**Fig. 1-figure supplement 2.**
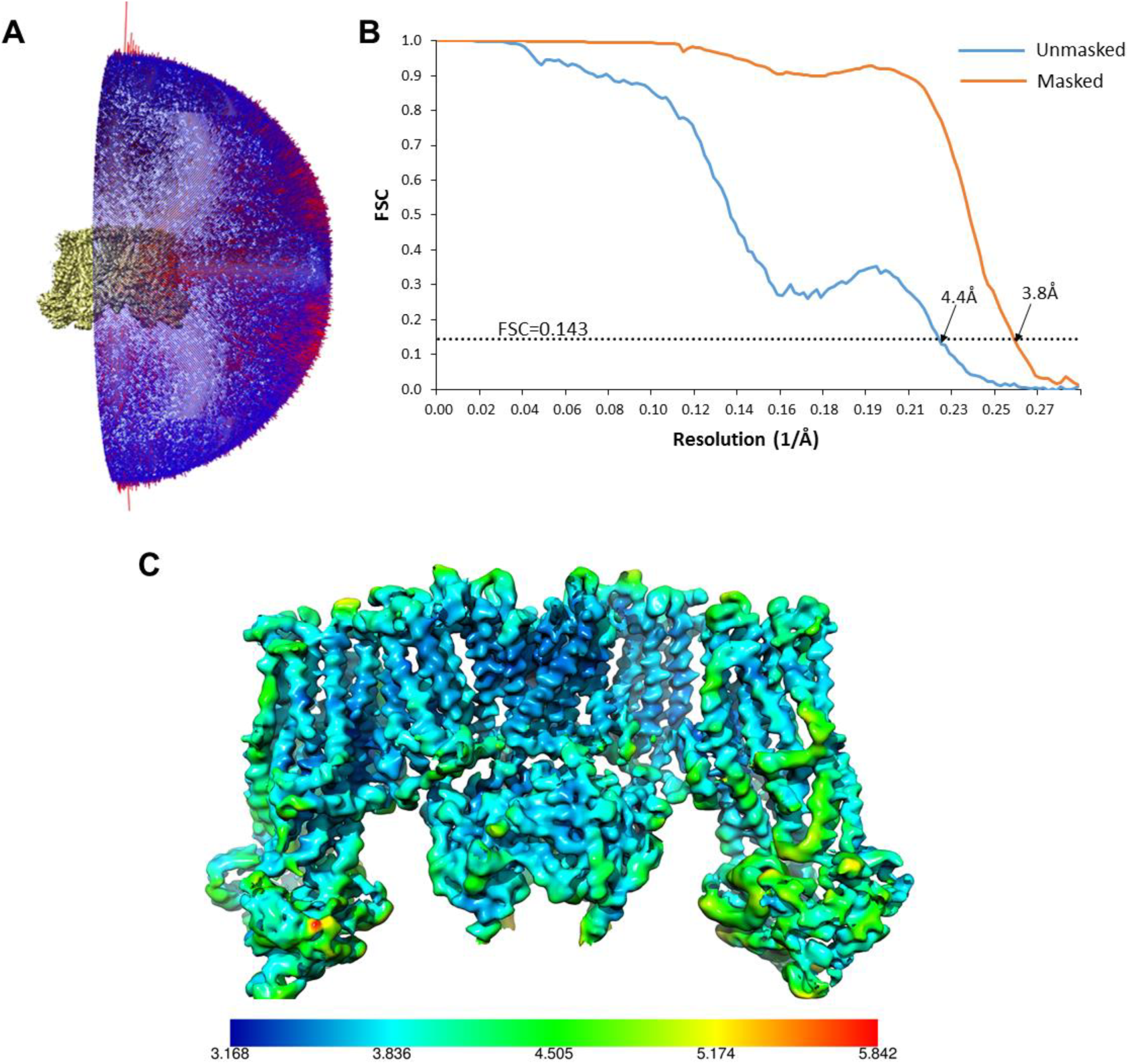
Cryo-EM density map analysis. **(A)** Euler angle distribution plot of all particles included in the calculation of the final map. **(B)** Fourier shell coefficient (FSC) curves of unmasked and masked whole complex. **(C)** The EM density map with colored local resolution estimation using Bsoft.

**Fig. 2-figure supplement 1.**
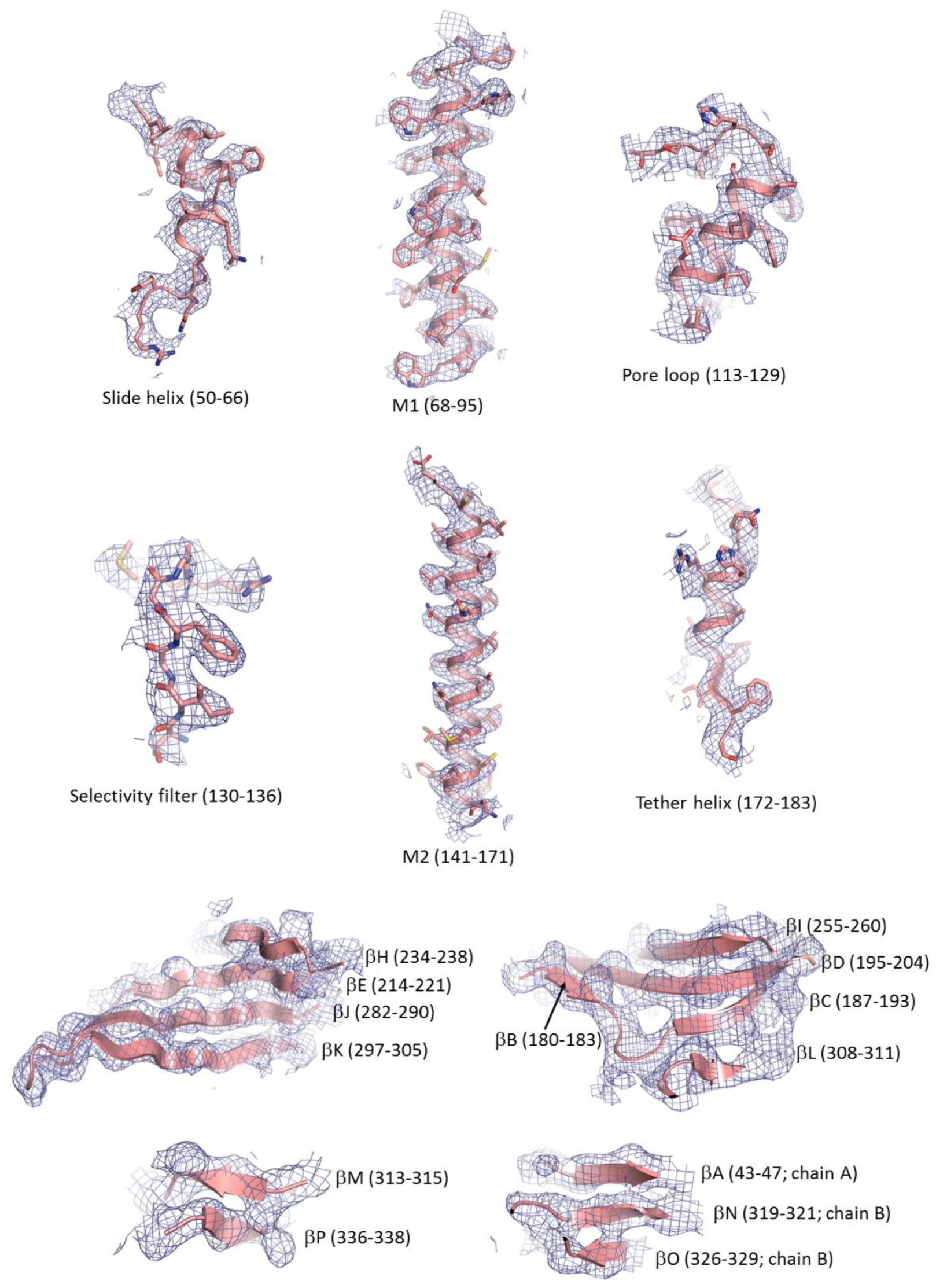
Electron density map of key structural features in Kir6.2. For the β-sheets only backbone is shown.

**Fig. 2-figure supplement 2.**
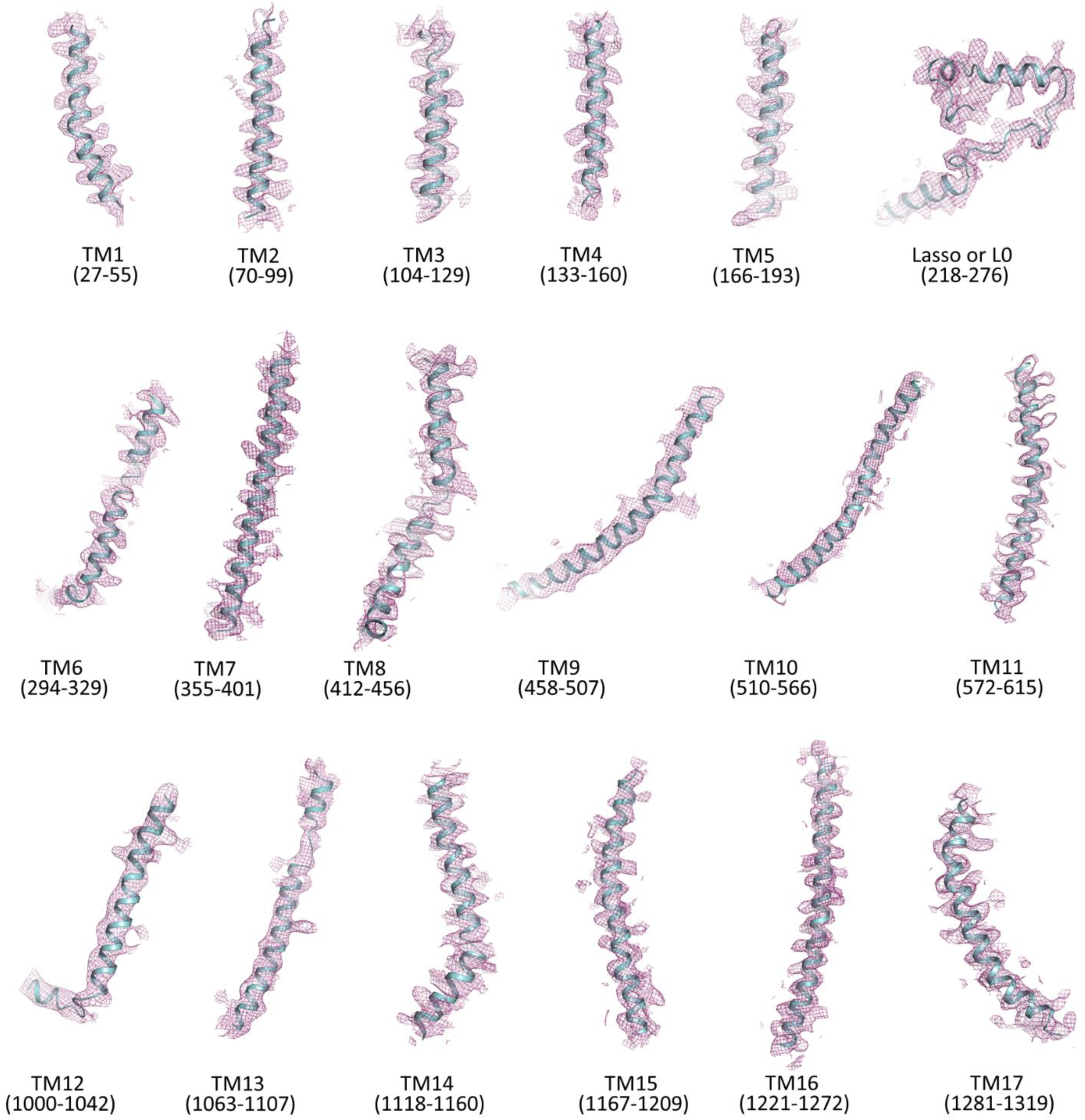
Electron density map of transmembrane helices and the lasso (L0) motif of SUR1. Only backbone is shown.

**Fig.4.-figure supplement 1.**
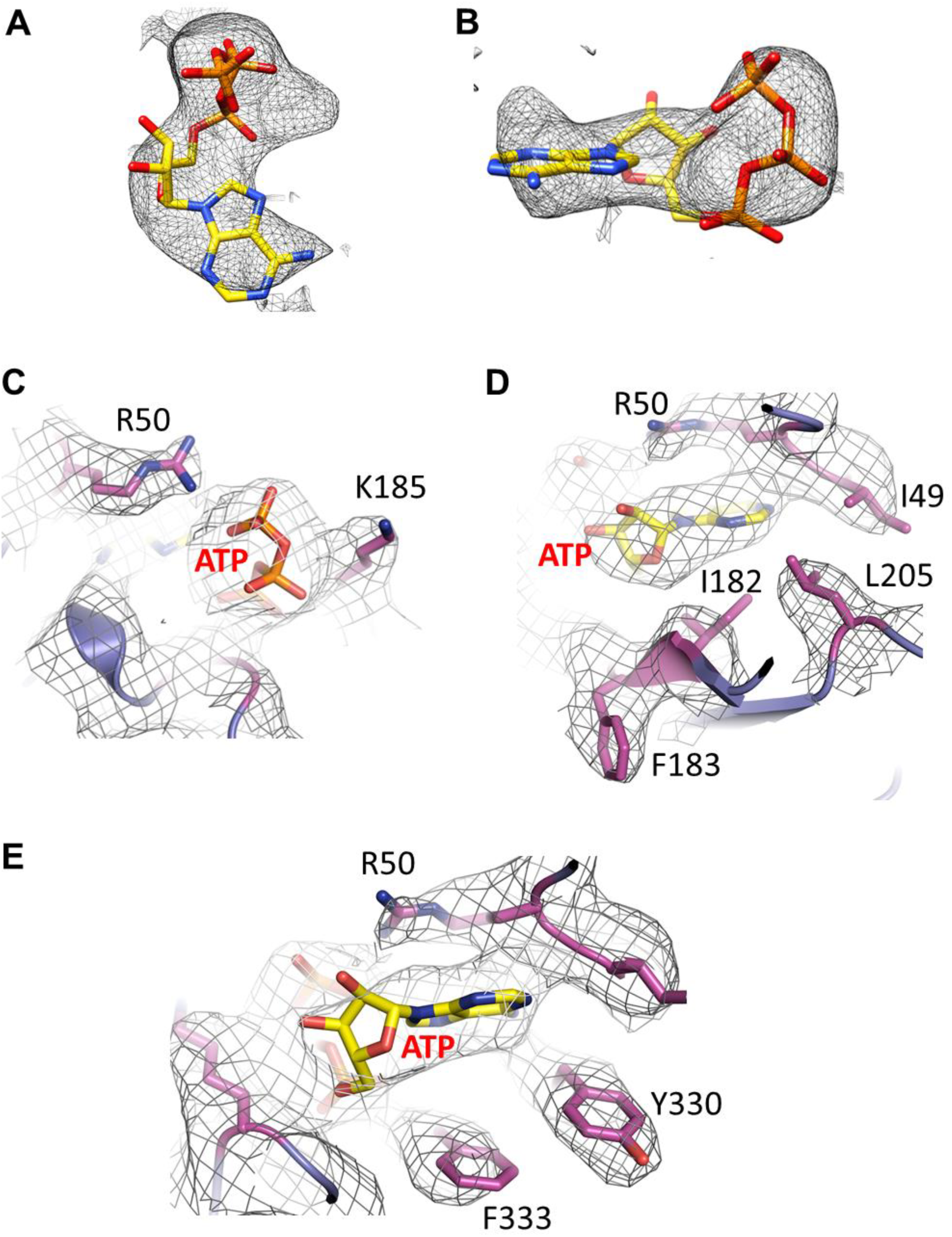
**(A and B)** Cryo-EM density for ATP, contoured to 3.5σ. **(C, D, E)** Cryo-EM density for residues surrounding ATP.

**Fig.6-figure supplement 1.**
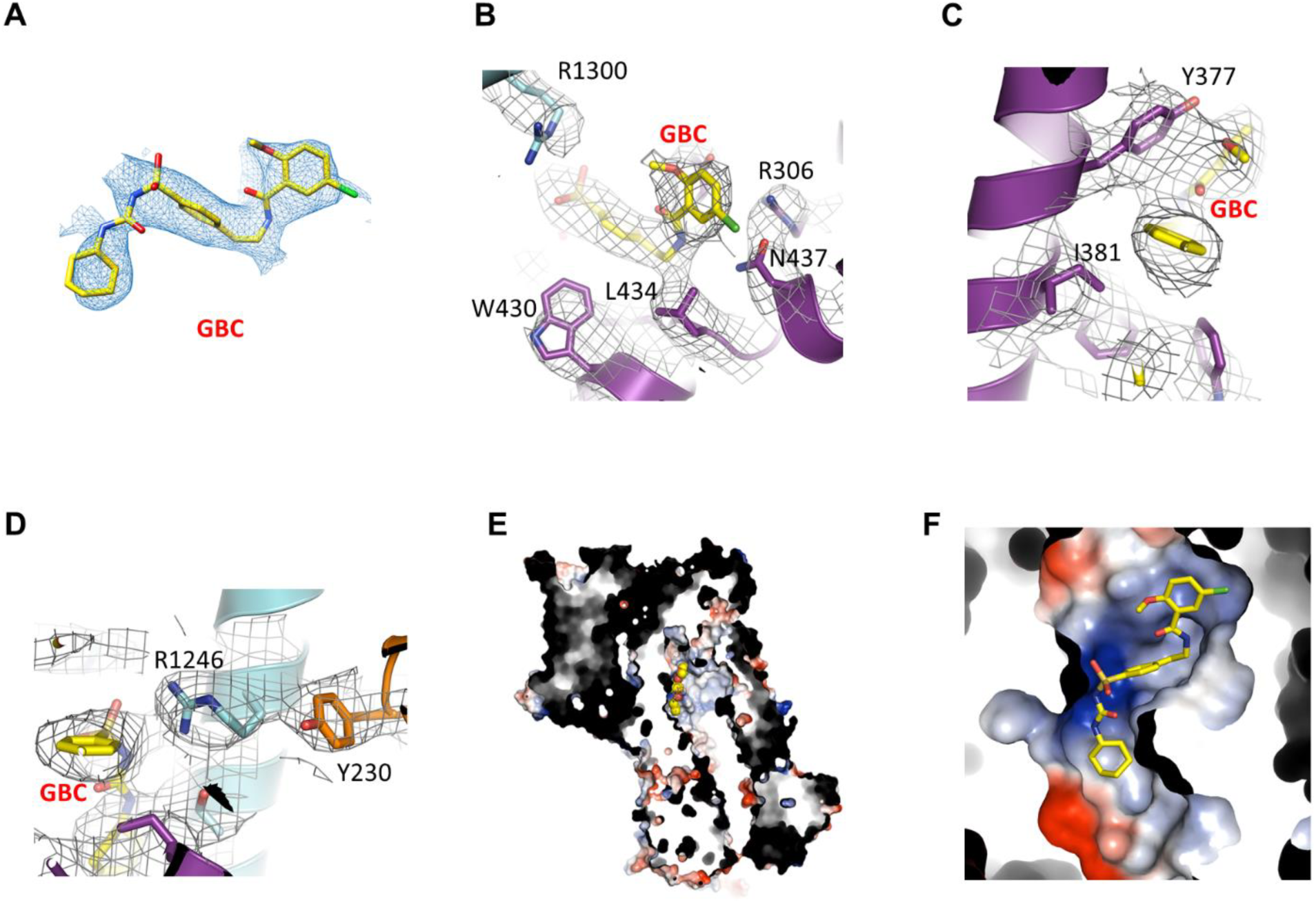
GBC binding site. **(A)** Cryo-EM density of GBC, contoured to 3σ. **(B, C, D)** Cryo-EM density of residues near GBC, contoured to 3.5σ. **(E)** Slice view of surface electrostatic potential, with GBC shown as spheres. Note how the binding pocket is formed by one end of the inner cleft between TMD1 and TMD2, abutting L0. **(F)** Close up view of GBC binding pocket. The basic portion comprises N1245, R1246, and R1300, while the acid end is formed by S1238 and D1193.

**Fig. 7-figure supplement 1.**
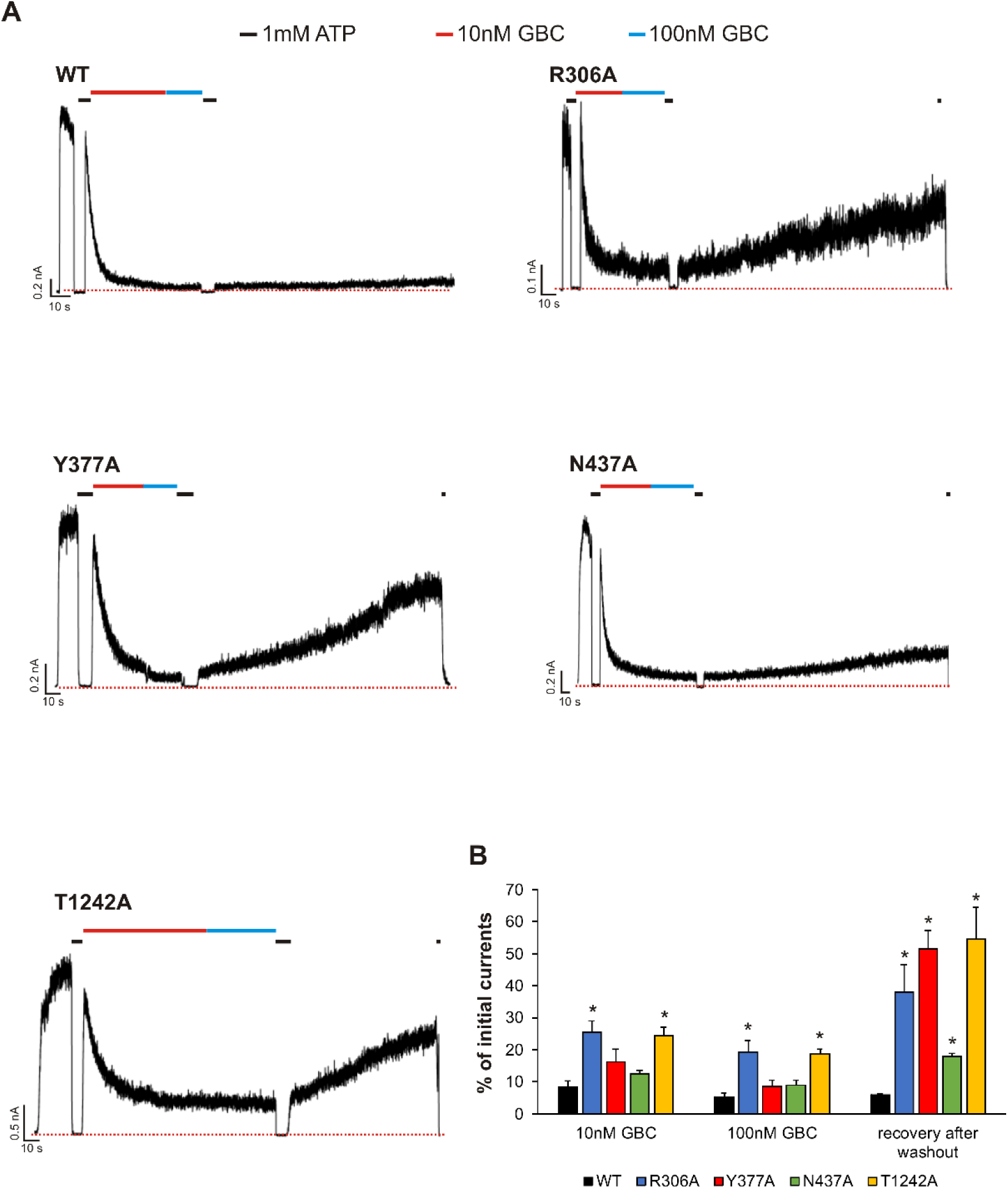
Functional testing of GBC binding residues by electrophysiology. **(A)** Examples of inside-out patch clamp recordings of WT and various mutant channels. Recordings were made at +50mV in symmetric K^+^ solutions and inward currents shown as upward deflections. **(B)** Quantification of residual currents (expressed as percent of initial currents observed in K-INT solution) after exposure to 10nM or 100nM GBC, with values taken when the currents reached a steady level. The value for “recovery after washout” was taken at ~100 seconds after the patch was returned to K-INT solution following a brief exposure to 1mM ATP (to check baseline). Each bar represents mean±s.e.m. of 3-6 patches. * *p*<0.05 by one-way ANOVA with Newman-Keuls *post hoc* test.

## References

Aguilar-Bryan, L., and J. Bryan. 1999. Molecular biology of adenosine triphosphate-sensitive potassium channels. Endocrine reviews. 20:101-135.

Aguilar-Bryan, L., and J. Bryan. 2008. Neonatal diabetes mellitus. Endocrine reviews. 29:265–291.

Antcliff, J.F., S. Haider, P. Proks, M.S. Sansom, and F.M. Ashcroft. 2005. Functional analysis of a structural model of the ATP-binding site of the KATP channel Kir6.2 subunit. The EMBO journal. 24:229–239.

Ashcroft, F.M. 2005. ATP-sensitive potassium channelopathies: focus on insulin secretion. The Journal of clinical investigation. 115:2047–2058.

Ashcroft, F.M. 2007. The Walter B. Cannon Physiology in Perspective Lecture, 2007. ATP-sensitive K+ channels and disease: from molecule to malady. American journal of physiology. Endocrinology and metabolism. 293:E880–889.

Ashfield, R., F.M. Gribble, S.J. Ashcroft, and F.M. Ashcroft. 1999. Identification of the high-affinity tolbutamide site on the SUR1 subunit of the K(ATP) channel. Diabetes. 48:1341–1347.

Brunger, A.T., P.D. Adams, G.M. Clore, W.L. DeLano, P. Gros, R.W. Grosse-Kunstleve, J.S. Jiang, J. Kuszewski, M. Nilges, N.S. Pannu, R.J. Read, L.M. Rice, T. Simonson, and G.L. Warren. 1998. Crystallography & NMR system: A new software suite for macromolecular structure determination. Acta crystallographica. Section D, Biological crystallography. 54:905–921.

Bryan, J., W.H. Vila-Carriles, G. Zhao, A.P. Babenko, and L. Aguilar-Bryan. 2004. Toward linking structure with function in ATP-sensitive K+ channels. Diabetes. 53 Suppl 3:S104–112.

Chapman, M.S., A. Trzynka, and B.K. Chapman. 2013. Atomic modeling of cryo-electron microscopy reconstructions--joint refinement of model and imaging parameters. Journal of structural biology. 182:10–21.

Chen, P.C., E.M. Olson, Q. Zhou, Y. Kryukova, H.M. Sampson, D.Y. Thomas, and S.L. Shyng. 2013a. Carbamazepine as a novel small molecule corrector of trafficking-impaired ATP-sensitive potassium channels identified in congenital hyperinsulinism. J. Biol. Chem. 288:20942–20954.

Chen, P.C., E.M. Olson, Q. Zhou, Y. Kryukova, H.M. Sampson, D.Y. Thomas, and S.L. Shyng. 2013b. Carbamazepine as a novel small molecule corrector of trafficking-impaired ATP-sensitive potassium channels identified in congenital hyperinsulinism. The Journal of biological chemistry. 288:20942–20954.

Cukras, C.A., I. Jeliazkova, and C.G. Nichols. 2002. The role of NH2-terminal positive charges in the activity of inward rectifier KATP channels. The Journal of general physiology. 120:437–446.

Devaraneni, P.K., G.M. Martin, E.M. Olson, Q. Zhou, and S.L. Shyng. 2015. Structurally distinct ligands rescue biogenesis defects of the KATP channel complex via a converging mechanism. The Journal of biological chemistry. 290:7980–7991.

Drain, P., L. Li, and J. Wang. 1998. KATP channel inhibition by ATP requires distinct functional domains of the cytoplasmic C terminus of the pore-forming subunit. Proceedings of the National Academy of Sciences of the United States of America. 95:13953–13958.

Gribble, F.M., and F. Reimann. 2003. Sulphonylurea action revisited: the post-cloning era. Diabetologia. 46:875–891.

Haider, S., J.F. Antcliff, P. Proks, M.S. Sansom, and F.M. Ashcroft. 2005. Focus on Kir6.2: a key component of the ATP-sensitive potassium channel. Journal of molecular and cellular cardiology. 38:927–936.

Hansen, S.B., X. Tao, and R. MacKinnon. 2011. Structural basis of PIP2 activation of the classical inward rectifier K+ channel Kir2.2. Nature. 477:495–498.

Hattori, M., and E. Gouaux. 2012. Molecular mechanism of ATP binding and ion channel activation in P2X receptors. Nature. 485:207–212.

Hohmeier, H.E., H. Mulder, G. Chen, R. Henkel-Rieger, M. Prentki, and C.B. Newgard. 2000. Isolation of INS-1-derived cell lines with robust ATP-sensitive K+ channel-dependent and -independent glucose-stimulated insulin secretion. Diabetes. 49:424–430.

Inagaki, N., T. Gonoi, J.P. Clement, C.Z. Wang, L. Aguilar-Bryan, J. Bryan, and S. Seino. 1996. A family of sulfonylurea receptors determines the pharmacological properties of ATP-sensitive K+ channels. Neuron. 16:1011–1017.

Inagaki, N., T. Gonoi, J.P.t. Clement, N. Namba, J. Inazawa, G. Gonzalez, L. Aguilar-Bryan, S. Seino, and J. Bryan. 1995. Reconstitution of IKATP: an inward rectifier subunit plus the sulfonylurea receptor. Science. 270:1166–1170.

Johnson, Z.L., and J. Chen. 2017. Structural Basis of Substrate Recognition by the Multidrug Resistance Protein MRP1. Cell. 168:1075–1085 e1079.

Kimanius, D., B.O. Forsberg, S.H. Scheres, and E. Lindahl. 2016. Accelerated cryo-EM structure determination with parallelisation using GPUs in RELION-2. eLife. 5.

Koster, J.C., M.A. Permutt, and C.G. Nichols. 2005. Diabetes and insulin secretion: the ATP-sensitive K+ channel (K ATP) connection. Diabetes. 54:3065–3072.

Koster, J.C., Q. Sha, and C.G. Nichols. 1999. Sulfonylurea and K(+)-channel opener sensitivity of K(ATP) channels. Functional coupling of Kir6.2 and SUR1 subunits. The Journal of general physiology. 114:203–213.

Kuhner, P., R. Prager, D. Stephan, U. Russ, M. Winkler, D. Ortiz, J. Bryan, and U. Quast. 2012. Importance of the Kir6.2 N-terminus for the interaction of glibenclamide and repaglinide with the pancreatic K(ATP) channel. Naunyn-Schmiedeberg's archives of pharmacology. 385:299–311.

Li, L., X. Geng, M. Yonkunas, A. Su, E. Densmore, P. Tang, and P. Drain. 2005. Ligand-dependent linkage of the ATP site to inhibition gate closure in the KATP channel. The Journal of general physiology. 126:285–299.

Li, N., J.X. Wu, D. Ding, J. Cheng, N. Gao, and L. Chen. 2017. Structure of a Pancreatic ATP-Sensitive Potassium Channel. Cell. 168:101–110 e110.

Lin, C.W., Y.W. Lin, F.F. Yan, J. Casey, M. Kochhar, E.B. Pratt, and S.L. Shyng. 2006. Kir6.2 mutations associated with neonatal diabetes reduce expression of ATP-sensitive K+ channels: implications in disease mechanism and sulfonylurea therapy. Diabetes. 55:1738–1746.

Lin, C.W., F. Yan, S. Shimamura, S. Barg, and S.L. Shyng. 2005. Membrane phosphoinositides control insulin secretion through their effects on ATP-sensitive K+ channel activity. Diabetes. 54:2852–2858.

Lin, Y.W., J.D. Bushman, F.F. Yan, S. Haidar, C. MacMullen, A. Ganguly, C.A. Stanley, and S.L. Shyng. 2008. Destabilization of ATP-sensitive potassium channel activity by novel KCNJ11 mutations identified in congenital hyperinsulinism. The Journal of biological chemistry. 283:9146–9156.

Liu, F., Z. Zhang, L. Csanady, D.C. Gadsby, and J. Chen. 2017. Molecular Structure of the Human CFTR Ion Channel. Cell. 169:85–95 e88.

Martin, G.M., C. Yoshioka, E.A. Rex, J.F. Fay, Q. Xie, M.R. Whorton, J.Z. Chen, and S.L. Shyng. 2017. Cryo-EM structure of the ATP-sensitive potassium channel illuminates mechanisms of assembly and gating. eLife. 6.

Nichols, C.G. 2006. KATP channels as molecular sensors of cellular metabolism. Nature. 440:470–476.

Notredame, C., D.G. Higgins, and J. Heringa. 2000. T-Coffee: A novel method for fast and accurate multiple sequence alignment. Journal of molecular biology. 302:205–217.

Payen, L., L. Delugin, A. Courtois, Y. Trinquart, A. Guillouzo, and O. Fardel. 2001. The sulphonylurea glibenclamide inhibits multidrug resistance protein (MRP1) activity in human lung cancer cells. British journal of pharmacology. 132:778–784.

Pettersen, E.F., T.D. Goddard, C.C. Huang, G.S. Couch, D.M. Greenblatt, E.C. Meng, and T.E. Ferrin. 2004. UCSF Chimera--a visualization system for exploratory research and analysis. Journal of computational chemistry. 25:1605–1612.

Pratt, E.B., F.F. Yan, J.W. Gay, C.A. Stanley, and S.L. Shyng. 2009. Sulfonylurea receptor 1 mutations that cause opposite insulin secretion defects with chemical chaperone exposure. The Journal of biological chemistry. 284:7951–7959.

Pratt, E.B., Q. Zhou, J.W. Gay, and S.L. Shyng. 2012. Engineered interaction between SUR1 and Kir6.2 that enhances ATP sensitivity in KATP channels. The Journal of general physiology. 140:175–187.

Proks, P., J.F. Antcliff, J. Lippiat, A.L. Gloyn, A.T. Hattersley, and F.M. Ashcroft. 2004. Molecular basis of Kir6.2 mutations associated with neonatal diabetes or neonatal diabetes plus neurological features. Proceedings of the National Academy of Sciences of the United States of America. 101:17539–17544.

Proks, P., F.M. Gribble, R. Adhikari, S.J. Tucker, and F.M. Ashcroft. 1999. Involvement of the N-terminus of Kir6.2 in the inhibition of the KATP channel by ATP. The Journal of physiology. 514 (Pt 1):19–25.

Proks, P., K. Shimomura, T.J. Craig, C.A. Girard, and F.M. Ashcroft. 2007. Mechanism of action of a sulphonylurea receptor SUR1 mutation (F132L) that causes DEND syndrome. Human molecular genetics. 16:2011–2019.

Reimann, F., S.J. Tucker, P. Proks, and F.M. Ashcroft. 1999. Involvement of the n-terminus of Kir6.2 in coupling to the sulphonylurea receptor. The Journal of physiology. 518 (Pt 2):325–336.

Rohou, A., and N. Grigorieff. 2015. CTFFIND4: Fast and accurate defocus estimation from electron micrographs. Journal of structural biology. 192:216–221.

Sagen, J.V., H. Raeder, E. Hathout, N. Shehadeh, K. Gudmundsson, H. Baevre, D. Abuelo, C. Phornphutkul, J. Molnes, G.I. Bell, A.L. Gloyn, A.T. Hattersley, A. Molven, O. Sovik, and P.R. Njolstad. 2004. Permanent neonatal diabetes due to mutations in KCNJ11 encoding Kir6.2: patient characteristics and initial response to sulfonylurea therapy. Diabetes. 53:2713–2718.

Schultz, B.D., A.D. DeRoos, C.J. Venglarik, A.K. Singh, R.A. Frizzell, and R.J. Bridges. 1996. Glibenclamide blockade of CFTR chloride channels. The American journal of physiology. 271:L192–200.

Smart, O.S., J.G. Neduvelil, X. Wang, B.A. Wallace, and M.S. Sansom. 1996. HOLE: a program for the analysis of the pore dimensions of ion channel structural models. Journal of molecular graphics. 14:354–360, 376.

Sola, D., L. Rossi, G.P. Schianca, P. Maffioli, M. Bigliocca, R. Mella, F. Corliano, G.P. Fra, E. Bartoli, and G. Derosa. 2015. Sulfonylureas and their use in clinical practice. Archives of medical science : AMS. 11:840–848.

Song, Y., F. DiMaio, R.Y. Wang, D. Kim, C. Miles, T. Brunette, J. Thompson, and D. Baker. 2013. High-resolution comparative modeling with RosettaCM. Structure. 21:1735–1742.

Tammaro, P., C. Girard, J. Molnes, P.R. Njolstad, and F.M. Ashcroft. 2005. Kir6.2 mutations causing neonatal diabetes provide new insights into Kir6.2-SUR1 interactions. The EMBO journal. 24:2318–2330.

Tucker, S.J., F.M. Gribble, P. Proks, S. Trapp, T.J. Ryder, T. Haug, F. Reimann, and F.M. Ashcroft. 1998. Molecular determinants of KATP channel inhibition by ATP. The EMBO journal. 17:3290–3296.

Tusnady, G.E., B. Sarkadi, I. Simon, and A. Varadi. 2006. Membrane topology of human ABC proteins. FEBS letters. 580:1017–1022.

Vila-Carriles, W.H., G. Zhao, and J. Bryan. 2007. Defining a binding pocket for sulfonylureas in ATP-sensitive potassium channels. FASEB journal : official publication of the Federation of American Societies for Experimental Biology. 21:18–25.

Voss, N.R., C.K. Yoshioka, M. Radermacher, C.S. Potter, and B. Carragher. 2009. DoG Picker and TiltPicker: software tools to facilitate particle selection in single particle electron microscopy. Journal of structural biology. 166:205–213.

Wilkens, S. 2015. Structure and mechanism of ABC transporters. F1000prime reports. 7:14.

Winkler, M., D. Stephan, S. Bieger, P. Kuhner, F. Wolff, and U. Quast. 2007. Testing the bipartite model of the sulfonylurea receptor binding site: binding of A-, B-, and A + B-site ligands. The Journal of pharmacology and experimental therapeutics. 322:701–708.

Yan, F.F., J. Casey, and S.L. Shyng. 2006. Sulfonylureas correct trafficking defects of disease-causing ATP-sensitive potassium channels by binding to the channel complex. The Journal of biological chemistry. 281:33403–33413.

Yan, F.F., Y.W. Lin, C. MacMullen, A. Ganguly, C.A. Stanley, and S.L. Shyng. 2007. Congenital hyperinsulinism associated ABCC8 mutations that cause defective trafficking of ATP-sensitive K+ channels: identification and rescue. Diabetes. 56:2339–2348.

Zerangue, N., B. Schwappach, Y.N. Jan, and L.Y. Jan. 1999. A new ER trafficking signal regulates the subunit stoichiometry of plasma membrane K(ATP) channels. Neuron. 22:537–548.

Zhang, Z., and J. Chen. 2016. Atomic Structure of the Cystic Fibrosis Transmembrane Conductance Regulator. Cell. 167:1586–1597 e1589.

Zheng, S.Q., E. Palovcak, J.P. Armache, K.A. Verba, Y. Cheng, and D.A. Agard. 2017. MotionCor2: anisotropic correction of beam-induced motion for improved cryo-electron microscopy. Nature methods. 14:331–332.

